# Targeting transcription factors through an IMiD independent zinc finger domain

**DOI:** 10.1101/2024.01.03.574032

**Authors:** Bee Hui Liu, Miao Liu, Sridhar Radhakrishnan, Chaitanya Kumar Jaladanki, Chong Gao, Jing Ping Tang, Kalpana Kumari, Mei Lin Go, Kim Anh L. Vu, Hyuk-Soo Seo, Kijun Song, Xi Tian, Li Feng, Justin L. Tan, Mahmoud A. Bassal, Haribabu Arthanari, Jun Qi, Sirano Dhe-Paganon, Hao Fan, Daniel G. Tenen, Li Chai

**Affiliations:** Cancer Science Institute of Singapore, Singapore 117599, Singapore; Department of Pathology, Brigham & Women’s Hospital, Harvard Medical School, Boston, MA 02115, USA; Bioinformatics Institute, Agency for Science, Technology and Research, Matrix 138671, Singapore; Department of Pharmacy, National University of Singapore; Department of Cancer Biology, Dana-Farber Cancer Institute, Boston, MA 02215, USA; Department of Biological Chemistry and Molecular Pharmacology, Harvard Medical School, Boston, MA 02115, USA; Department of Medicine, Harvard Medical School, Boston, MA 02115, USA; Institute of Cancer Research, Shenzhen Bay Laboratory, Shenzhen 518132, China; Harvard Stem Cell Institute, Harvard Medical School, Boston, MA 02115, USA

## Abstract

Immunomodulatory imide drugs (IMiDs) degrade specific C2H2 zinc finger degrons in transcription factors, making them effective against certain cancers. SALL4, a cancer driver, contains seven C2H2 zinc fingers in four clusters, including an IMiD degron in zinc finger cluster two (ZFC2). Surprisingly, IMiDs do not inhibit growth of SALL4 expressing cancer cells. To overcome this limit, we focused on a non-IMiD degron, SALL4 zinc finger cluster four (ZFC4). By combining AlphaFold and the ZFC4-DNA crystal structure, we identified a potential ZFC4 drug pocket. Utilizing an *in silico* docking algorithm and cell viability assays, we screened chemical libraries and discovered SH6, which selectively targets SALL4-expressing cancer cells. Mechanistic studies revealed that SH6 degrades SALL4 protein through the CUL4A/CRBN pathway, while deletion of ZFC4 abolished this activity. Moreover, SH6 led to significant 62% tumor growth inhibition of SALL4+ xenografts in vivo and demonstrated good bioavailability in pharmacokinetic studies. In summary, these studies represent a new approach for IMiD independent drug discovery targeting C2H2 transcription factors in cancer.

**Graphical Abstract:** 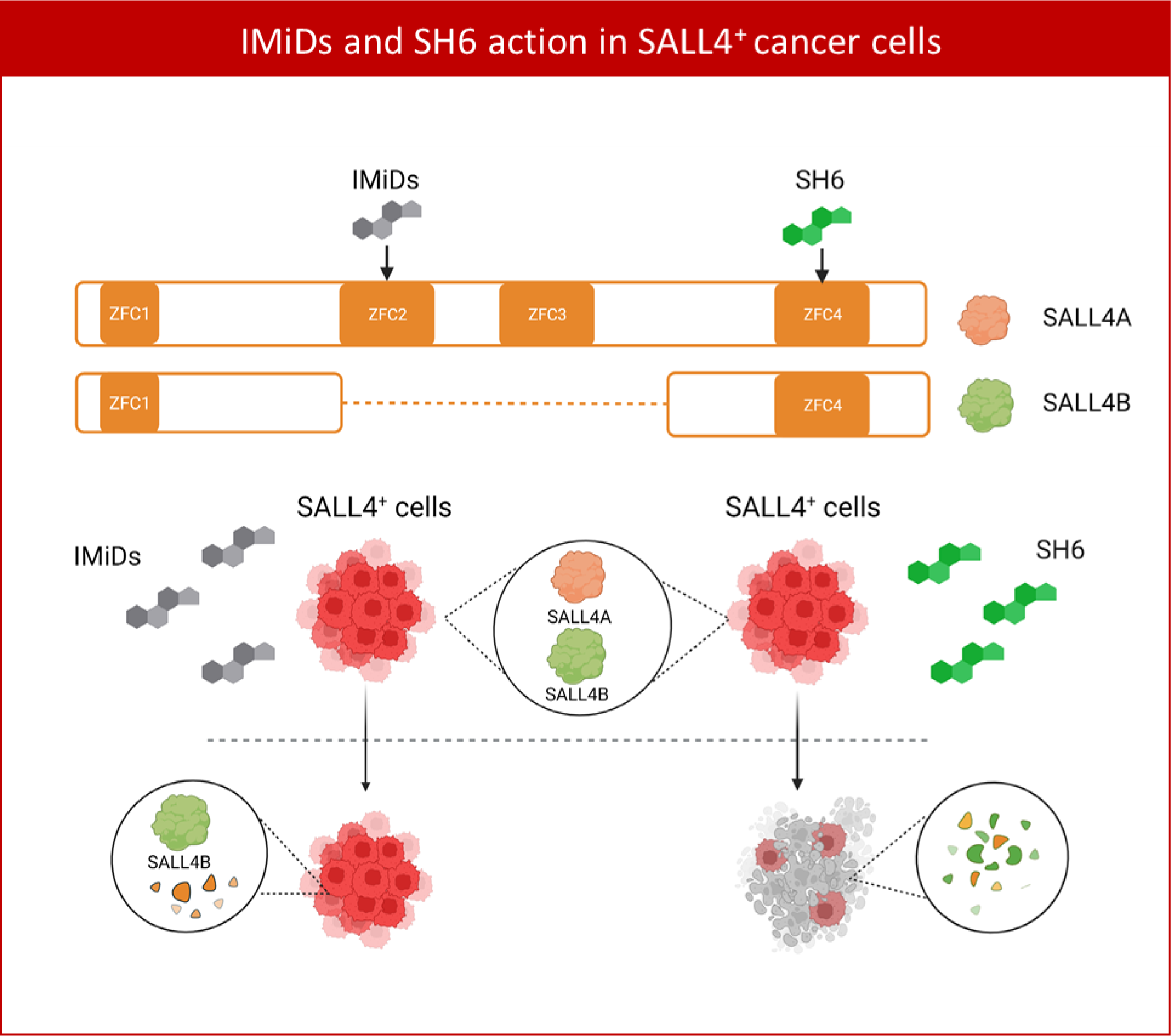

## Introduction

Transcription factors (TFs) have been historically viewed to be “undruggable”(Bushweller, 2019), with a number of attempts with mixed results(Grembecka, Graf et al., 2007, Masso-Valles & Soucek, 2020, Rego, He et al., 2000, Verhoeven, Tilborghs et al., 2020). Recent breakthroughs and successes in induced protein degradation are among the most exciting new frontiers for development of novel cancer drugs against TFs. Examples of such approaches include clinically approved Immunomodulatory imide drugs (IMiDs), or by proteolysis-targeting chimera (PROTAC) technology.(Filippakopoulos, Qi et al., 2010, Jan, Sperling et al., 2021) The IMiD drugs Thalidomide, Lenalidomide, and Pomalidomide bind to Cereblon (CRBN), a substrate receptor of the CUL4-RBX1-DDB1-CRBN (CRL4CRBN) E3 ubiquitin ligase, altering its substrate selectivity to recruit and degrade a specific set of proteins(Lu, Middleton et al., 2014). Beside the known targeted proteins of IMiDs such as Ikaros (IKZF1), Aiolos (IKZF3), Casein kinase 1 alpha (CK1α), GSPT1, and SALL4, a set of Cys2-His2 (C2H2) zinc fingers has also been identified to be degraded by Thalidomide analogs(Sievers, Petzold et al., 2018).

SALL4 is a zinc finger transcription factor that is important for many cancers, most notably hepatocellular carcinoma (HCC)(Yong, Gao et al., 2013), lung cancer(Kobayashi, Kuribayashi et al., 2011), ovarian cancer(Yang, Xie et al., 2016), breast cancer(Kobayashi, Kuribayshi et al., 2011), myelodysplastic syndrome (MDS)(Liu, Kwon et al., 2022), and acute myeloid leukemia (AML)(Li, Yang et al., 2013, Ma, Cui et al., 2006). While we have recently described a peptidomimetic molecule that has the ability to inhibit SALL4 function in HCC cell lines and xenograft mouse models(Liu, Jobichen et al., 2018), no small molecule targeted therapy against SALL4 in cancer is currently available.

SALL4 has 2 main isoforms. While SALL4A, the larger isoform, is translated from the full-length SALL4 gene, SALL4B, the smaller isoform, is a spliced variant lacking the region of exon 2 encoding zinc finger clusters 2 and 3 (ZFC2 and ZFC3), while both isoforms contain ZFC4, the DNA binding domain(Kong, Bassal et al., 2021). The crystal structure of SALL4 ZFC2, which is not present in SALL4B, has been shown to bind thalidomide(Furihata, Yamanaka et al., 2020) and pomalidomide(Matyskiela, Clayton et al., 2020) together with CRBN. We recently observed that although IMiDs are capable of degrading SALL4A through ZFC2, they do not target the SALL4B isoform. As a result, IMiDs demonstrate no effectiveness in viability of cancer cell lines that express both SALL4A and SALL4B(Kong et al., 2021, Vu, Kumari et al., 2023). To target SALL4+ cancer cells, we need to degrade both isoforms, therefore, we focused on targeting ZFC4, which is shared by both SALL4A and SALL4B.

Previously, we and others characterized the consensus DNA sequence that SALL4 ZFC4 binds(Kong et al., 2021, Pantier, Chhatbar et al., 2021). Here, we determined the crystal structure of SALL4’s ZFC4/DNA complex, and aided by the recent AlphaFold(Bryant, Pozzati et al., 2022, Flower & Hurley, 2021, Jumper & Hassabis, 2022, Porta-Pardo, Ruiz-Serra et al., 2022, Varadi, Anyango et al., 2022) program, which predicts protein structure folds with high accuracy, we identified a potential binding pocket in SALL4 ZFC4. After identifying a potential druggable pocket in ZFC4, we screened chemical libraries to isolate a lead small molecule SH6, which could degrade SALL4 protein and inhibit the growth of SALL4+ cancer cells in culture *and in vivo*.

## Results

### Combining AlphaFold, crystal structure, and molecular docking to screen known and novel drug libraries for SALL4 ZFC4 targeting compounds

As described above, SALL4 has two isoforms: SALL4A and SALL4B (Fig. 1A). We(Kong et al., 2021) and others(Pantier et al., 2021) recently independently found that the DNA binding domain of SALL4 is located in the fourth C2H2 zinc finger cluster (ZFC4) and that it prefers a consensus SALL4 DNA binding motif(Kong et al., 2021, Pantier et al., 2021). Unlike ZFC2, which is only present in SALL4A, ZFC4 is a common domain shared by SALL4A and SALL4B. To further determine the structural basis of the interaction between SALL4 and DNA, we co-crystallized two molecules of SALL4 ZFC4 (amino acids 864-929) with a 12-mer blunt-end duplex DNA containing the SALL4 TATTA DNA binding motif (CGAAATATTAGC)(Kong et al., 2021). The crystal structure was solved at a resolution of 2.1Å in a space group with an asymmetric unit that contained one molecule of the duplex DNA and two molecules of SALL4 ZFC4 (Fig. 1B). The statistics are shown in Table EV1. While the first zinc finger of the second SALL4 molecule (ZFC4a’) did not generate electron density, the remaining three zinc fingers (ZFC4a, ZFC4b and ZFC4b’) generated high-quality electron density, and bound to the duplex in canonical fashion, namely by inserting their helical elements into the major groove of DNA (Fig. EV1). Unlike the first finger ZFC4a, which provided only backbone interactions, the second finger (ZFC4b) interacted specifically with the amide side chain of Asn912 (N912), via the adenines at positions D4 and D5 of one strand and C10 of the other strand (Fig. EV1). Side chains of the second zinc finger of both molecules also provided four and five additional interactions with the DNA backbone, respectively, including Lys896-Y4, Arg905-Y3, Lys910-X4, Lys914-X5, and His916-Y2. Recently, the crystal structure of human SALL4 ZFC4 (856–930)(Ru, Koga et al., 2022) and murine SALL4 ZFC4 (870–940)(Watson, Pantier et al., 2023) were published. Consistent with the report from Ru et al(Ru et al., 2022), our crystallographic findings of human SALL4 ZFC4 (864-929) bound to its consensus DNA binding target revealed an asymmetric unit that contained one molecule of the duplex DNA and two molecules of ZFC4. Both our crystal and the human ZFC4(856-930) crystal(Ru et al., 2022) demonstrated that only one of the two zinc-fingers of the second ZFC4 molecule is bound, suggesting the non-DNA binding zinc finger could be involved in other unknown function(s).

**Figure 1.**
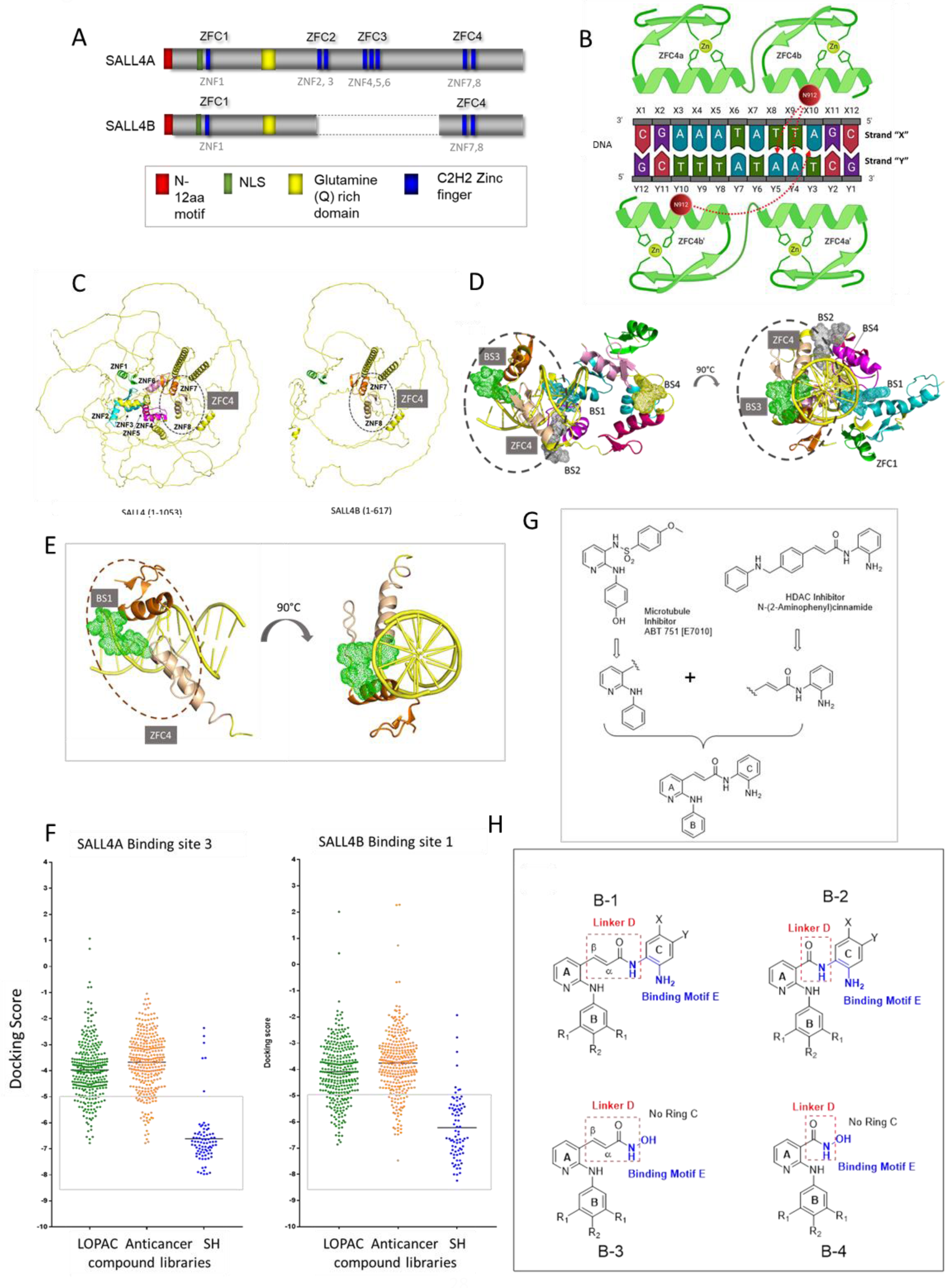
Identification of a novel ZFC4 binding site. **A)** Schematic representation of SALL4 protein isoforms. Dark blue rectangles = zinc fingers. SALL4A consists of four C2H2 zinc finger clusters (ZFC1, ZFC2, ZFC3, & ZFC4), while SALL4B, an internally spliced isoform, contains only ZFC1 and ZFC4. ZFC1 consists of Zinc Finger (ZNF) 1, ZFC2 consists of ZNF2&3, ZFC3 consists of ZNF4, 5, and 6, while ZFC4 consists of ZNF7&8. **B)** Ligand Interaction Diagram of the SALL4 ZFC4-DNA complex is shown schematically. Key SALL4 residues (N912) are labelled within red circles. Zinc ions are shown as small green circles. The duplex DNA strands are labelled arbitrarily with letters Y and X and numbered sequentially. Key interactions are shown as dotted red arrows. **C)** AlphaFold2 prediction of SALL4A and SALL4B. Left panel: ZNF1 (Green), ZNF2 (Cyan), ZNF3 (Teal), ZNF4 (magenta), ZNF5 (hot pink), ZNF6 (pink), ZNF7 (orange), and ZNF8 (wheat). Right panel: SALL4B, highlighted ZNF4 (ZNF7 (orange) and ZNF8 (wheat)). **D)** and **E)** Binding site prediction of SALL4 using SALL4 models from **(A)** and the ZFC4-DNA structure. Top view and side view of predicted binding sites on the SALL4A **(D)** and SALL4B **(E**) are shown. **F)** small molecule docking screen against the common ZFC4 binding site in SALL4A (BS3) and SALL4B (BS1). Green = LOPAC1280 library, Orange= FDA approved anti-cancer library, Blue=SH library. **G)** The SH series of compounds are based on a hybrid design that incorporates the N-phenylpyridin-2-amino motif (rings A, B) of E7010 to the HDAC N-(2-aminophenyl) acrylamide moiety (ring C). **H)** The following scaffolds are found in the SH series: B-1 in SH1-SH7, SH9-SH15, AH17-SH23, SH25-SH31, SH33-SH39 are characterized by an acrylamide linker D; B-2 is representative of SH41-SH47, SH49-SH55, SH57-SH63, SH65-SH71, SH73-SH79 in which linker E is an amide; B-3 are arylamides in which ring C is replaced by OH (SH8,SH16,SH24,SH32,SH40); B-4 are amides in which ring C is replaced by OH (SH48,SH56,SH64,SH72,SH80).

We next tested the possibility of targeting SALL4 by molecular docking based on our knowledge of this newly characterized DNA binding structure. As the available ZFC4 crystal structure only encompasses a small region of the full-length SALL4 proteins, and there is no available SALL4B model in UniProt currently, we therefore used AlphaFold2(Jumper, Evans et al., 2021, Jumper & Hassabis, 2022) to generate the 3D structural models of both isoforms (Fig. 1C). We first generated 5 models of SALL4A, selected the best scored model (pLDDT score), and compared it to the SALL4A model in UniProt. The comparison between our best scored model of SALL4A and the SALL4A model available in UniProt revealed a low Root Mean Square Deviation (RMSD) of 0.81 Å, further supporting our model. Subsequently, we generated SALL4B models using AlphaFold2 and selected the best scored model. After superimposing our ZFC4 crystal data with the AlphaFold models, we observed a strong congruence between them, with a root mean square deviation (RMSD) of 0.90 Å. Since the AlphaFold models were generated without any ligands, we predicted binding sites on the SALL4 isoforms using SiteMap(Schrödinger, 2020d). Next, we transferred the DNA from our crystal structure to the AlphaFold models by superimposition and predicted binding sites. For SALL4A, four binding sites were predicted: binding site 1 (BS1) in ZFC2 & 3; binding site 2 (BS2) in ZFC3 & 4; binding site 3 (BS3) within ZFC4; and binding site 4 (BS4) in ZFC3 (Fig. 1D). For SALL4B, only one binding site was predicted in ZFC4 that was identical to binding site 3 in SALL4A (SALL4B BS1, Fig. 1E). Not only is this site the only site shared between SALL4A and B, but it is also a site with potential clefts between the protein and DNA macromolecules that could be relevant for small molecule binding, including a water-filled albeit hydrophobic pocket of approximately 600 A^2^ formed at the intersection of the two zinc-finger molecules and the DNA. Therefore, this common site was chosen for subsequent work.

Chemical libraries with a total of 676 compounds from known (Sigma Aldrich LOPAC and Selleck Chem Anticancer Library) and novel (in house SH Library) small molecule libraries were docked against SALL4A BS3 and SALL4B BS1 (Fig. 1F). Using an arbitrary cut off of -5.0 for docking scores, we shortlisted 119 compounds for further study (Tables 1 and 2). Interestingly, compounds from the in-house CSI-SH library surpassed the other libraries in docking to both binding sites in SALL4A and SALL4B. This library consists of 80 dual motif molecules (Table 2) bearing pharmacophores in which the anti-microtubule N-phenylpyridin-2-amino motif (rings A, B) of the microtubule inhibiting agent E7010 were linked to the HDAC benzamide moiety (ring C) of HDAC 1/2 benzamide zinc binders via an acrylamide linker D (Fig. 1G). The following scaffolds are found in the SH series: B-1 are characterized by an acrylamide linker D; B-2 with amide linker D; B-3 are acrylamides in which ring C is replaced by OH; and B-4 are amides in which ring C is replaced by OH (Fig. 1H).

**Table 1.** Selected compound from LOPAC and Anticancer libraries based on docking scores on ZFC4 binding site.

| LOPAC1280 |  |  |
| --- | --- | --- |
| Name | SALL4A-DNA Binding site 3 | SALL4B-DNA Binding site 1 |
| Ancitabine hydrochloride | -6.78 | -6.77 |
| Indinavir sulfate salt hydrate | -6.63 | -6.51 |
| (±)-Norepinephrine (+)bitartrate | -6.54 | -6.53 |
| L(±)-Norepinephrine (+)bitartrate | -6.54 | -6.53 |
| (±)-HA-966 | -6.32 | -6.02 |
| N6-2-(4-Aminophenyl)ethyladenosine | -6.13 | -6.69 |
| SKF-89145 hydrobromide | -6.05 | -5.58 |
| Cytosine-1-beta-D-arabinofuranoside | -5.99 | -6.23 |
| 1-Aminobenzotriazole | -5.84 | -5.15 |
| Adenosine | -5.82 | -6.08 |
| 2-Chloroadenosine | -5.74 | -5.66 |
| D-Cycloserine | -5.62 | -5.47 |
| DL-Cycloserine | -5.62 | -5.47 |
| 5-azacytidine | -5.59 | -6.21 |
| Aminophylline ethylenediamine | -5.54 | -5.64 |
| L-Cycloserine | -5.49 | -5.41 |
| Gemcitabine hydrochloride | -5.49 | -6.16 |
| Acyclovir | -5.40 | -5.25 |
| 4-Aminopyridine | -5.39 | -5.28 |
| Apomorphine hydrochloride hemihydrate | -5.27 | -5.42 |
| Allopurinol | -5.26 | -6.04 |
| Indirubin-3'-oxime | -5.24 | -5.44 |
| GR79236X | -5.23 | -5.74 |
| Acetazolamide | -5.21 | -6.11 |
| Pyrocatechol | -5.20 | -6.41 |
| Alfuzosin hydrochloride | -5.18 | -5.90 |
| (±)-Baclofen | -5.16 | -5.56 |
| 4-Aminobenzamidine dihydrochloride | -5.14 | -5.31 |
| Gabaculine hydrochloride | -5.09 | -5.17 |
| HEMADO | -5.00 | -5.98 |
| 3'-Azido-3'-deoxythymidine | -5.00 | -5.57 |
| FDA approved anti-cancer compounds |  |  |
| Gemcitabine HCl (Gemzar) | -6.78 | -6.27 |
| Cytarabine | -6.73 | -6.48 |
| Canagliflozin | -6.63 | -7.47 |
| Dorzolamide HCl | -6.43 | -5.25 |
| Gemcitabine | -6.31 | -6.22 |
| Decitabine | -6.08 | -5.44 |
| Entinostat | -5.92 | -5.96 |
| Regorafenib | -5.92 | -5.78 |
| Fludarabine | -5.88 | -6.26 |
| Floxuridine | -5.82 | -6.47 |
| Quercetin | -5.81 | -5.65 |
| Veliparib | -5.80 | -5.42 |
| Nelarabine | -5.76 | -6.41 |
| Cladribine | -5.68 | -6.23 |
| Clofarabine | -5.67 | -6.34 |
| R406 (free base) | -5.60 | -5.72 |
| Ftorafur | -5.24 | -5.57 |
| Streptozotocin | -5.13 | -5.68 |
| (-)-Epigallocatechin gallate | -5.05 | -5.52 |

**Table 2.**
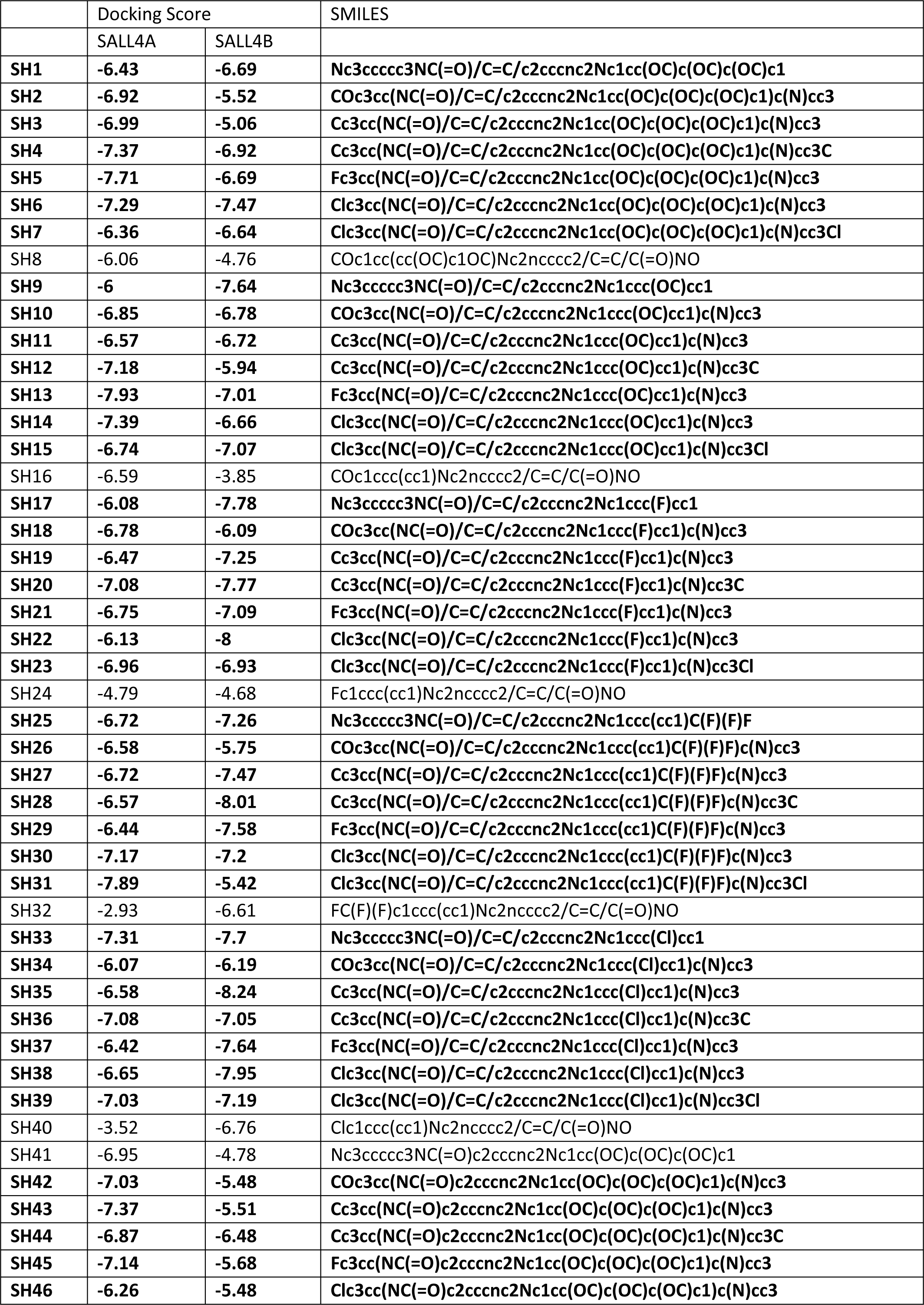
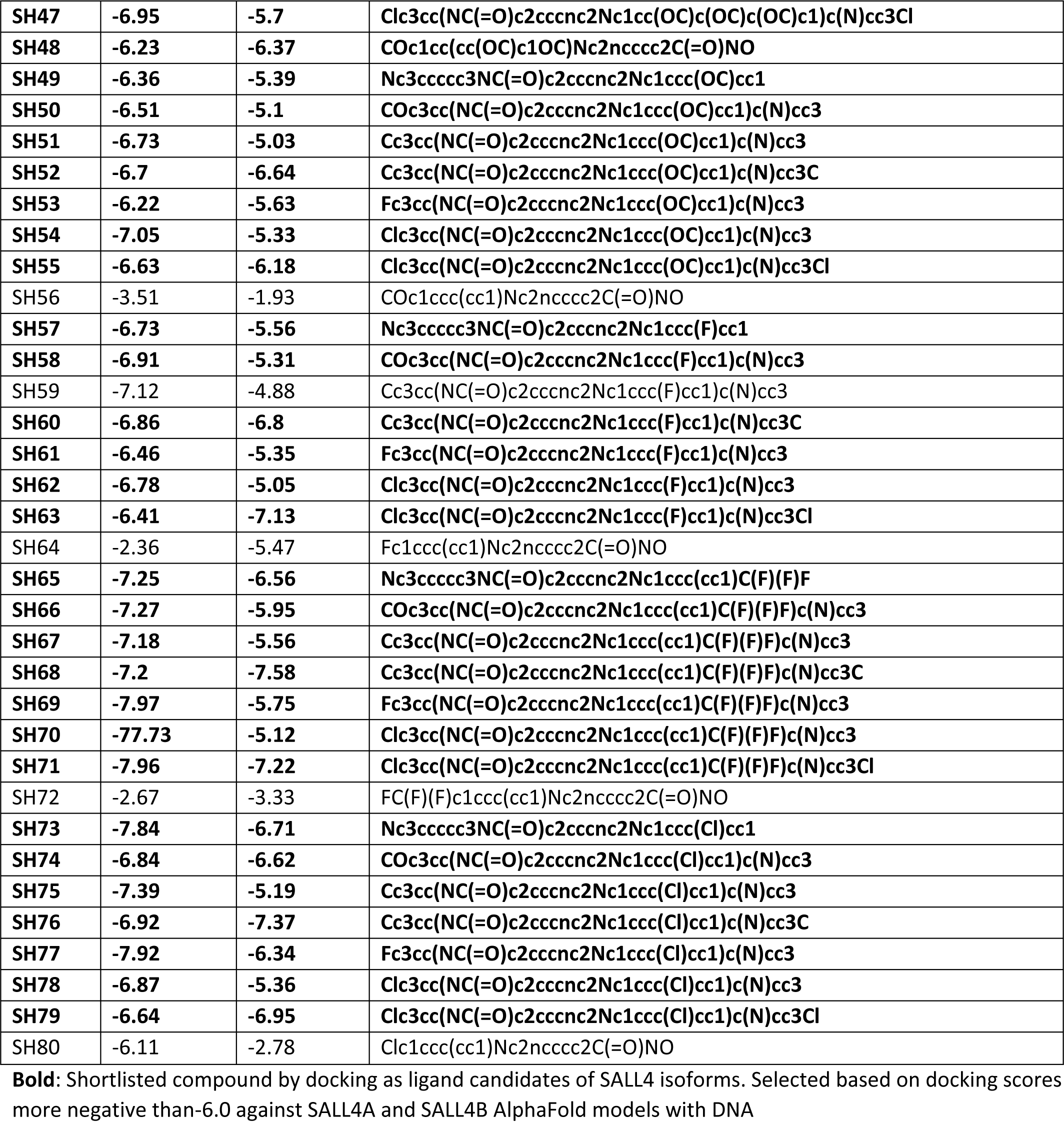
Docking scores of SH library on ZFC4 binding site.

### SH6 is a bona fide SALL4 targeting compound

Cell-based screening was next employed to evaluate the selected compounds in targeting SALL4 expressing malignant cells. In this system, a pair of liver cancer cell lines with high (SNU398) and negligible (SNU387) endogenous expression of SALL4, respectively, were used. Cells were incubated with 117 compounds (two compounds not available for testing) at 1μM for 72 h and cell viability assessed (Fig. 2A). Three compounds (SH2, SH6, SH7) displayed selective killing against SALL4 high cells. These three compounds, as well as the rest of SH library, were further selected for dose escalation studies to deduce the EC_50_ in SNU387, SNU398, and isogenic cell lines overexpressing SALL4A or SALL4B (SNU387-TgSALL4A and SNU387-TgSALL4B)(Tan, Li et al., 2019) (Fig. 2, Table 2, and Table EV2). The EC_50_ of SH2, SH6, and SH7 in SNU398 high cells were 0.4μM, 0.5μM, and 0.3μM, respectively. In contrast, the EC_50_ values of the same compounds were 50-100 fold higher on SALL4 low cells. Interestingly, the test compounds were consistently more potent (by approximately 3-fold) against the SALL4B high isoform as compared to the SALL4A high isoform. To determine if these compounds could target other SALL4 high malignant cells, we tested SH2, SH6, and SH7 against three lung cancer cell lines with high (SALL4^Hi^) or low (SALL4^Lo^) levels of SALL4 (Fig. 2C). A similar trend was again observed, with the test compounds displaying greater potencies against SALL4 expressing lung cancer cell lines. Taken together, the results support the notion that the test compounds preferentially targeted malignant cells that express high levels of SALL4.

**Figure 2.**
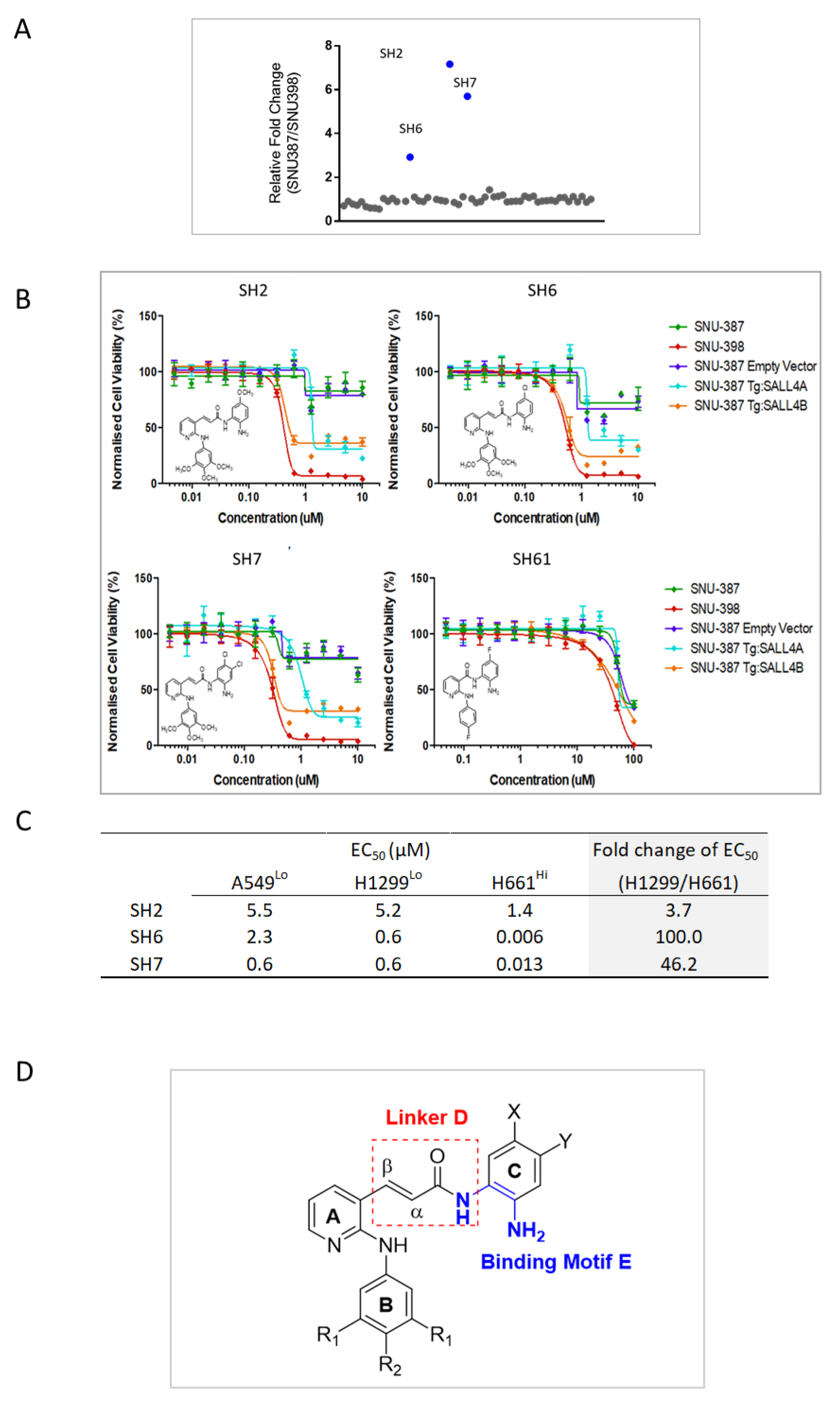
Identification of a novel class of SALL4 inhibitor. **A)** Single dose phenotypic screen using SALL4 high (SNU398) vs low cell lines (SNU387) for 117 selected compounds from Fig 1f, Table 1 and Table 2; two of the compounds are not available for testing. **B)** SH library compounds were tested for EC50 in cell viability assays. Isogenic cell lines specifically expressing SALL4A (SNU387-TgSALL4A) or SALL4B (SNU387-TgSALL4B) were tested alongside SNU398 and SNU387 cells. SH2, SH6, SH7, and SH61 are shown in the representative graphs. **C)** SH2, 6, and 7 were utilized in dose dependent phenotypic screens using the lung cancer cell line H661 (SALL4 high) and H1299 and A549 (SALL4 low). The result was compiled into a table in (c). **D)** Schematic representation of three key structure-activity correlations in the SH library. A detailed description can be found in Fig. EV3. Each data point in (B) and (C) was perform in n=3 independent cell cultures. IC50 of compounds were determined using nonlinear regression fitting, with error bars represents s.d. of mean.

Next, we compared the molecular properties of SH2, SH6, and SH7 (Fig. EV2A) using the cheminformatics tools Molinspiration (https://www.molinspiration.com/). It was found that lipophilicity or hydrophobicity increases as SH7>SH6>SH2, and that SH6 is slightly more hydrophobic than SH2, with lesser solubility but higher permeability. We then determined the PAMPA permeability of SH6 and SH7; the effective permeability Pe of SH6 and SH7 were 66.13 x 10^-6^cm/sec and 41.16 x 10^-6^cm/sec, respectively (Fig. EV2A) compared to control compounds (highly permeable carbamazine Pe = 65.22 x 10^-6^ cm/sec, and poorly permeable antipyrine Pe =1.28 x 10 ^-6^cm/sec, Fig. EV2B determined under similar conditions). SH6 and SH7 have good permeability profiles, in keeping with their lipophilic character (mlog P 4.05 and 4.66, respectively). Due to its superiority in overall scores, we selected SH6 as the lead compound for further study.

Furthermore, three key structure-activity correlations were noted for the SH library (Fig. 2D). First, the acrylamide linker is an indispensable feature for activity. Second, the phenylene diamine ring C in the HDAC component of the hybrid design is needed for potent cell-based activities. Third, the sterically bulky and lipophilic trimethoxy substituted ring B is required to establish strong interactions. Detailed structural activity studies can be found in Fig.EV3.

### SH6 degrades SALL4B resulting in on-target cancer cell killing through the CUL4A-CRBN pathway

To understand the mechanism of SH6, we first probed SALL4A and SALL4B levels in SNU398 cells that have been treated with SH6 or Pomalidomide. While Pomalidomide treatment led to a notable decrease in SALL4A protein, SH6 treatment led to decreases of both SALL4A and SALL4B isoforms (Fig. 3A). Intriguingly, in spite of its ability to degrade SALL4A, the IMiD Lenalidomide failed to suppress the viability of SALL4 high SNU398 cells. In contrast, SH6 is able to selectively target SALL4 high cells (Fig. 3B). We have recently found that depletion of SALL4B is required for targeting SALL4-mediated tumorigenesis(Vu et al., 2023). To further examine the effect of IMiDs and SH6 in degrading SALL4B, SALL4B was fused to NanoLuc(Lu et al., 2014) and stably transfected into H1299 cells. Cells were then treated with SH6 or Lenalidomide for 24hr, followed by measurement of emitted luciferase as an indicator of SALL4B protein levels (Fig. 3C). The result demonstrated that SH6 treated cells emitted the least amount of luminescence as compared to Lenalidomide treated cells, indicating degradation of SALL4B by SH6 and not by Lenalidomide. Taken together, the results indicate that degradation of SALL4B by SH6 induced cell death, which could not be achieved by IMiDs in SALL4 expressing cancer cells.

**Figure 3.**
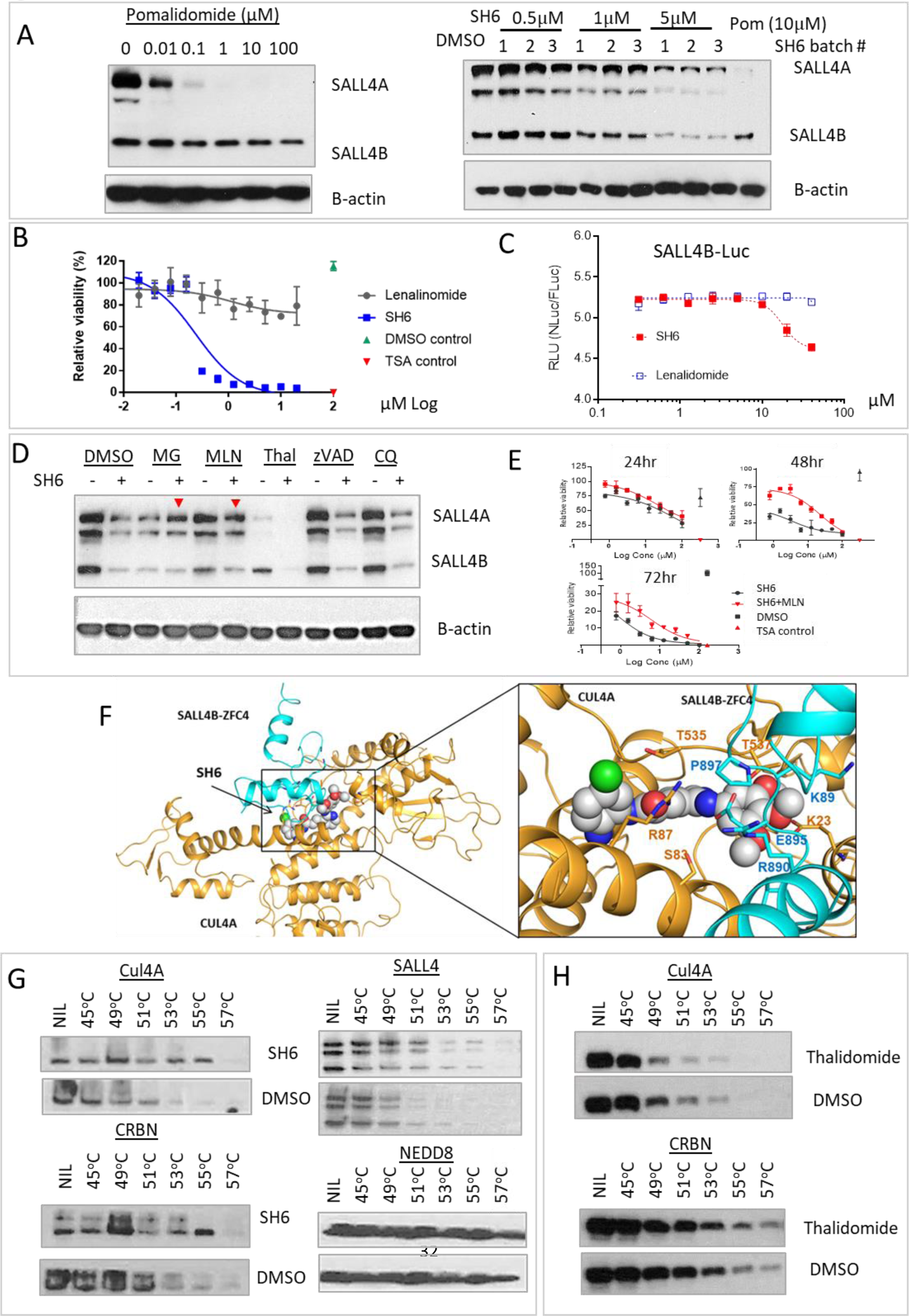
IMiD independent SH6 mediated CUL4A-CRBN degradation of SALL4. **A)** SNU398 cells were treated with Pomalidomide or SH6 at the indicated concentrations for 24 hours. Cells were harvested for western blot analysis. **B)** Cells were treated with SH6 or lenalidomide for 72 hours and examined for their viability. SH6 decreases cell viability while Lenalidomide does not show activity on these cells. **C)** Dual-luciferase reporter assay. SALL4B was fused to the NanoLuc dual luciferase vector (pLL3.7-EF1a-IRES-Gateway-nluc-2xHA-IRES2-fluc-hCL1-P2A-Puro) and stably transfected into SALL4+ H1299 lung cancer cells. Cells were treated with SH6 or lenalidomide for 24 hours and assayed for luminescence. DMSO normalized NLuc/FLuc data was presented as the mean ± s.d. of n = 3 biologically independent samples. **D)** DMSO, MLN4924 (MLN), MG132 (MG), Thalidomide (Thal), zVAD-FMK (zVAD), or Chloroquine (CQ) were added for four hours prior to SH6 treatment, followed by western blot analysis 24hrs later. MLN4924 and MG132 demonstrated prominent rescue of SH6 degradation of SALL4 (red arrows). **E)** MLN4924 rescue of SH6 induced cell killing. SNU398 cells were incubated with SH6 or SH6 in combination with MLN4924 (SH6+MN) for 24hr, 48hr, or 72hr. Cell viability was measured and data are presented as mean ± s.d. of n = 3 biologically independent samples. **F)** The docking pose of SH6 at the predicted SALL4B ZFC4-CUL4A interface. Cyan=SALL4B ZFC4; Orange= CUL4A **G)** Cells were incubated with SH6 for 4 hours prior to CETSA analysis of the Cullin pathway. In-cell CETSA was performed as described in Materials and Methods. Cell lysates were separated by SDS-PAGE and analyzed by Western blotting. In-cell CETSA demonstrated that SH6 prolongs thermal stability for CUL4A, CRBN, and SALL4, in comparison to DMSO. **H)** Cells treated with Thalidomide or DMSO underwent in-cell CETSA, revealing that Thalidomide stabilizes CRBN but not CUL4A. Each data point in (B) and (C) was perform in n=3 independent cell cultures. Inhibition curves were determined using nonlinear regression fitting, with error bars represents s.d. of mean.

Previous studies have reported that SALL4 is an IMiD-dependent CRL4^CRBN^ substrate(Donovan, An et al., 2018, Filippakopoulos et al., 2010). This led us to ask which pathway would be utilized under IMiD independent, SH6-mediated SALL4 degradation. To test this, SNU398 cells were treated with either DMSO, the proteasome inhibitor MG132, the NEDD8-activating enzyme (NAE) inhibitor MLN4924, the IMiD thalidomide, a pan caspase inhibitor zVAD-FMK, or a lysosomal inhibitor chloroquine (CQ), prior to SH6 treatment. Cell lysates were harvested after 24 hours of SH6 treatment, followed by western blot analysis (Fig. 3D). Interestingly, we noted that SH6 induced degradation of SALL4 was rescued by MG132, and more significantly by MLN4924, but not by zVAD or chloroquine. The result indicates that the SH6-SALL4 degradation pathway involves NEDD-8 mediated E3 ligase proteasomal degradation. When neddylation of cullin was blocked by MLN4924, SALL4 protein was rescued. Furthermore, when SNU398 cells were co-treated with MLN4924 and SH6, and cell viability monitored over time (24h, 48h, 72h), we observed a clear rescue profile at the latter two time points, which was noticeably absent in cells treated with SH6 alone under similar conditions (Fig. 3E). Taken together, these results implicate the involvement of an activated cullin pathway for SALL4 degradation.

We then took a computation-guided model prediction approach to determine if CRBN-SALL4 interacts with CUL4A. Using existing crystal structures of the SALL4 ZFC2-CRBN-DDB1 complex as reference point, the crystal structures of the SALL4-CRBN-DDB1 complex (PDB ID: 6UML) and the DDB1-CUL4A-Rbx1 complex (PDB ID: 2HYE) were superimposed (with the predicted complex renamed as SALL4-CRL4^CRBN^) (Fig. EV4). When full length SALL4 was superimposed over this SALL4-CRL4^CRBN^ complex, we observed that ZFC3 and ZFC4 clash with CUL4A near the RBX1 domain, indicating that SALL4 and CUL4A might interact with each other. Next, protein-protein docking analysis was performed with CUL4A (extracted from PDB ID: 2HYE), SALL4A ZFC 2 to 4, and SALL4B ZFC4 of SALL4B. Thereafter, we superimposed the SALL4-CUL4A complexes with the SALL4-CRL4^CRBN^ complex (defined above). We identified a potential binding site that appear in both ZFC2-4 of the SALL4A-CUL4A complex, and the ZFC4 of SALL4B-CUL4A complex (Fig. 3F), coinciding with BS3 of SALLA and BS1 of SALL4B identified previously (Fig. 1D and 1E). We docked SH6, SH61 (a negative control), as well as three thalidomide derivatives against this binding site (Table EV3). SH6 outperformed the other five compounds in the docking score. With its docking pose wedged between ZFC4 of SALL4A/B and CUL4A, we hypothesized that SH6 may also bind to CUL4A (Fig. 3F).

Next, we evaluated this model through a cell-based target engagement by SH6 using the cellular thermal shift assay (CETSA)(Jafari, Almqvist et al., 2014, Martinez Molina, Jafari et al., 2013). Briefly, when a small molecule binds a protein, it stabilizes the protein. Consequently, unfolding of the protein slows down at elevated temperatures. Here, we employed the in-cell CETSA to investigate the binding of SH6 to several components (NEDD8, CUL4, and CRBN) of the cullin pathway, and SALL4 itself. SNU398 cells were incubated with SH6 or DMSO for four hours prior to harvesting and lysis for CETSA analysis (Fig. 3G). We found that while the NEDD8 unfolding profile was not altered by SH6, the presence of SH6 led to higher thermal stability (less unfolding) of CUL4, CRBN, and SALL4 (Fig. 3G), indicating that these proteins are bound to SH6. A similar CETSA was performed using the IMiD Thalidomide, and only unfolding of CRBN, and not CUL4A, was delayed in the presence of the IMiD compound (Fig. 3H). Taken together, our results indicate that even though SH6 is structurally different from IMiDs, it mediates SALL4 degradation via the CUL4A^CRBN^ pathway. Unlike IMiDs, which do not directly target SALL4B, SH6 binds to both SALL4 and CUL4 to induce the degradation of SALL4B, resulting in death of cancer cells that express SALL4.

### SH6 is a bona fide ZFC4 degrader

Our earlier findings highlight the significance of drug-ZFC4 binding in facilitating the degradation of SALL4B by SH6 to induce cancer cell death. To further examine the importance of ZFC4 in SH6 mediated SALL4B degradation, we generated 293T cells that either expressed SALL4B (SALL4B^WT^), or a SALL4B mutant lacking ZFC4 (SALL4B^-ZFC4^). The cells were subsequently treated with SH6, Lenalidomide, or DMSO for 24hrs, before harvested for western blot analysis (Fig 4A). The results indicated that SH6 effectively degraded wild type SALL4B protein, but failed to degrade SALL4B in the absence of ZFC4. In contrast, Lenalidomide did not induce degradation of either the wild type SALL4B or the SALL4 mutant. Top

**Figure 4.**
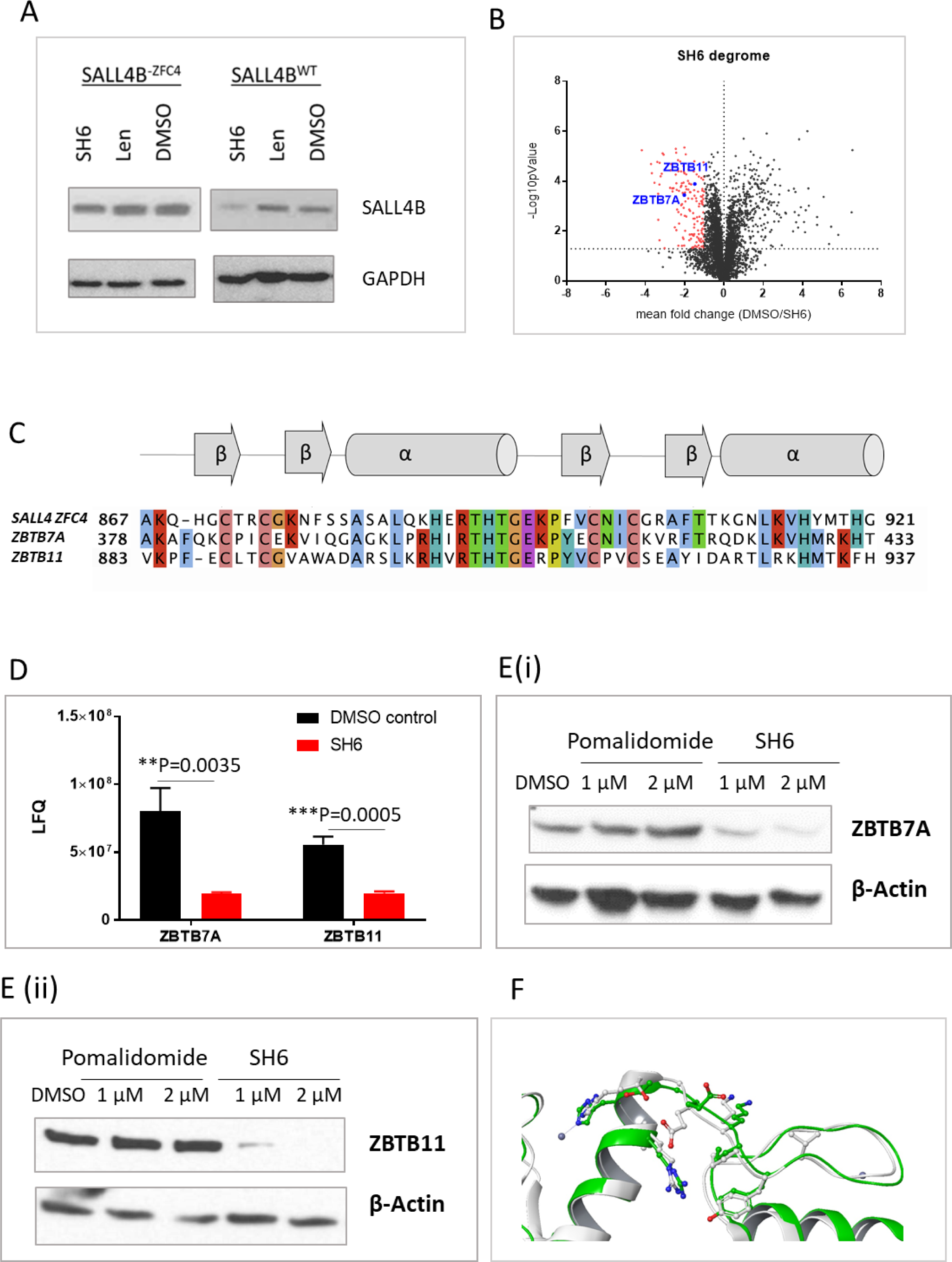
SH6 is a ZFC4 degrader. **A)** 293T cells were stably transfected with pFUW-SALL4B wild type (SALL4B^WT^), or pFUW-SALL4B with deletion of ZFC4 (SALL4B^-ZFC4^). Cells were treated with SH6, Lenalidomide (Len), or DMSO for 24 hours before harvested for western blot analysis. SH6 mediated degradation of SALL4B was abolished in the absence of ZFC4, while Lenalidomide did not affect SALL4B levels in SALL4B^-ZFC4^ or SALL4B^WT^cells. **B)** Degrome analysis revealed two additional ZFC4 containing proteins. A volcano plot from the mass spectrometry data demonstrates the change of the cellular proteins from the SH6 vs DMSO treatment. Down regulated proteins with twofold change and p-value of 0.05 were highlighted in red. In brief, SNU398 was grown treated with SH6 (50uM) or DMSO for 24hr. **C)** Sequence alignment of the SALL4 ZFC4 domain with ZBTB7A and ZBTB11. α-helices (α) and β-strands (β) are indicated according to data on crystal structures of SALL4 ZFC4. The colour scheme used for residue conservation is based on side chain properties. Blue=hydrophobic (A, I, L, M, F, W, V); Red=positive charge (K, R); Magenta= negative (D, E); Green= polar (S, T, N, Q); Pink= cysteine; Orange= glycine; Yellow= proline; Cyan= aromatic (Y, H). **D)** Expression levels of ZBTB7A and ZBTB11 in DMSO/ SH6 treated samples analysed by mass spectrometry. Data represent the mean ± s.d. (n=3). **E)** SNU398 cells were treated with DMSO, Pomalidomide, or SH6 for 24 hours, harvested, and analysed by western blot. Blots were probed with ZBTB7A (**E(i)**) or ZBTB11 (**E(ii)**) antibodies. **F)** The ZBTB7A crystal structure (PDB 7N5W; Green) was overlaid with that of SALL4 ZFC4 (Red) and analysed with PyMol. Overlapped regions are shown.

We next conducted mass spectrometry analysis on the SH6 degrome to search for additional ZFC4 targets of SH6. To do this, we treated SNU398 with SH6 for 24 hours and then performed mass spectrometry. Total proteome data was analysed using LFQ intensity values in a log2 scale. Proteins with 2-fold down regulation compared to DMSO, and with P values smaller than 0.05, were defined as down-regulated proteins (Fig.4B; Table EV4). Sequence alignments were performed on these down-regulated targets, compared to that of the sequence of SALL4 ZFC4, using Clustal Omega(Sievers & Higgins, 2018, Sievers, Wilm et al., 2011). We identified two C2H2 zinc finger-containing proteins, ZBTB7A and ZBTB11, out of the 153 down-regulated proteins, which showed high sequence similarity with SALL4 ZFC4. (Fig. 4C). Upon examination of the SH6 degrome, we found that ZBTB7A and ZBTB11 were down-regulated 4.1 and 2.8 times, respectively (Fig. 4D). When cells were treated with Pomalidomide or SH6, SH6 was noted to degrade both proteins starting at 1µM, whereas Pomalidomide did not cause degradation of ZBTB7A or ZBTB11 at the tested concentrations (Fig. 4E). Subsequently, we superimposed the crystal structure of ZBTB7A(Ren, Horton et al., 2023) (PDB ID: 7N5W) with the crystal structure of SALL4 ZFC4 and found that the two crystals superimposed at the ZF1-ZF2 region of ZBTB7A (Fig. 4F). Based on our findings, we can conclude that SH6 is a genuine IMiD independent ZFC4 degrader.

### SH6 inhibits tumor growth in mouse xenografts with a desirable pharmacokinetic profile

To determine selective cytotoxicity SH6, it was used to treat two immortalized liver cell lines derived from normal liver (THLE2 and THLE3) (Fig. 5A). We found that SH6 selectively kills SALL4 expressing liver cancer cells with EC_50_ values of 0.3μM to 1.3 μM (Table EV2), as compared to values exceeding 25 μM against the two immortalized nontransformed THLE2 and THLE3 liver cell lines.

**Figure 5.**
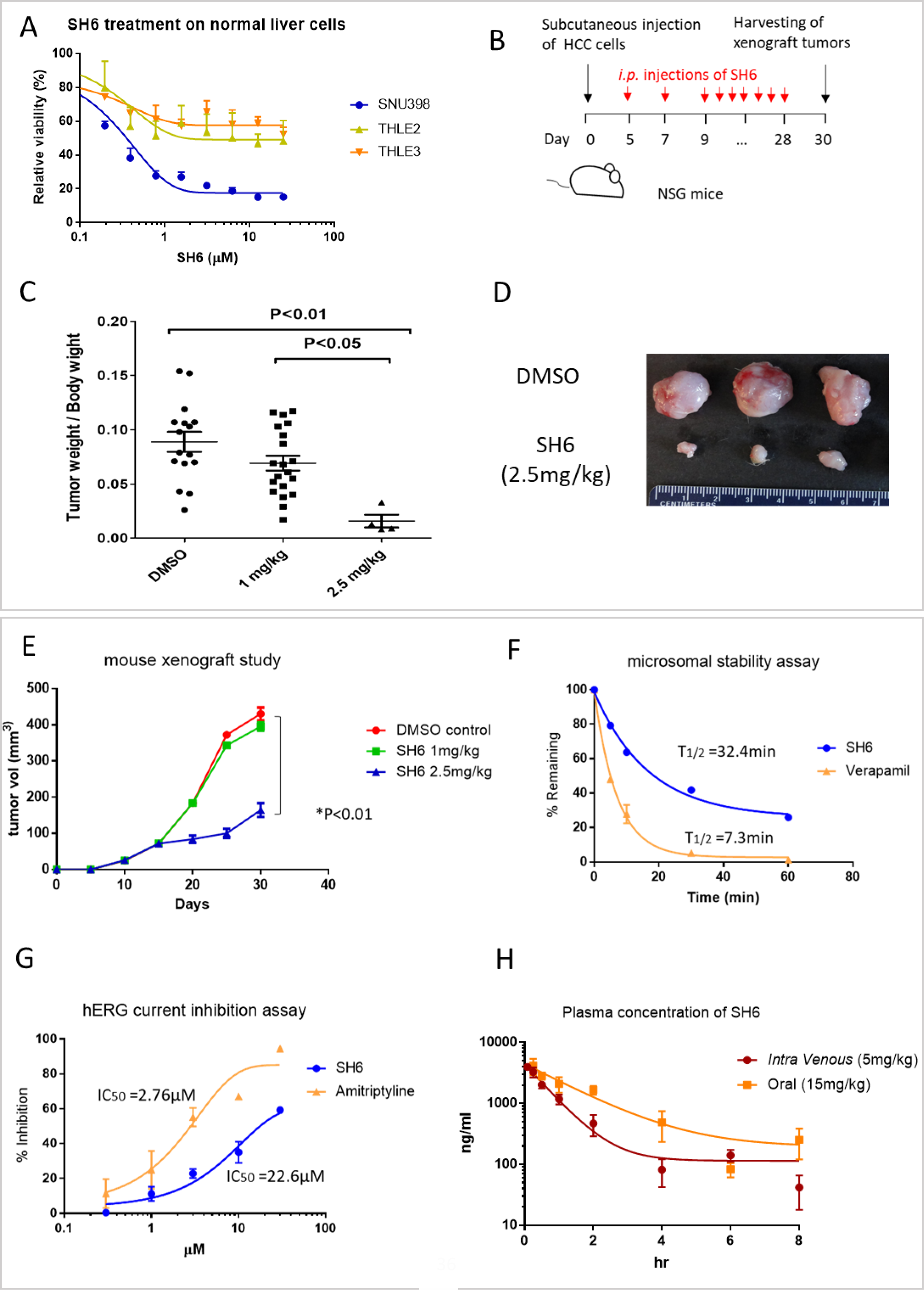
SH6 leads to tumor regression and demonstrates a satisfactory pharmacokinetic profile. **A)** SH6 specifically inhibits cell growth of SNU398 HCC but not non-transformed liver cell lines THLE-2 and THLE-3. **B)** Design of the mouse xenograft experiments. NSG mice were transplanted with 2x10^6^ SNU398 cells, and the tumors were allowed to form a palpable mass. SH6 or DMSO were administered every other day from Day 5 to Day 28, and tumors were harvested on Day 30. **C)** Tumors of the harvested xenografts at Day 30, with tumor weight at end point over mouse body weight. **D)** Size of three representative tumors at 30 days comparing DMSO versus SH6 treatment. **E)** Tumor growth was traced from Day 0 to Day 30 for DMSO, SH6 at 1mg/kg, and SH6 at 2.5mg/kg. **F)** Microsomal stability assay with 1μM of SH6 or Verapamil. The percentage of the remaining compound (% Remaining, Y axis) was analyzed with LC-MS/MS. **G)** Concentration response of hERG inhibition by SH6, with amitriptyline as the control. **H)** Single dose pharmacokinetic study of SH6 via intravenous injection (5mg/kg) or oral gavage (15mg/kg) on Swiss albino mice (n=3). The plasma concentration of the compound was measured with LC-MS/MS. Each data point in (A), (E), (F), (G) and (H) was perform in n=3 independent cell cultures. IC50 of compounds in (A), (F) and (H) were determined using nonlinear regression fitting, with error bars represents s.d. of mean. *P* values were calculated by two-tailed Student’s t-tests

To further test the anti-tumor potential of SH6, SALL4-high SNU398 cells were implanted subcutaneously into the flanks of NOD/SCID/Gamma mice (NSG), which were then randomly grouped for SH6 or DMSO treatment (Fig. 5B). Tumors in the DMSO control group (n=16) progressively increased in size over a period of 30 days. In contrast, SH6 at 2.5mg/kg (n=4) induced a strong therapeutic effect (p<0.01) with a tumor growth inhibition of 62%, compared with DMSO controls, or SH6 at a lower dose of 1 mg/kg (n=16) (p<0.01 and p<0.05, respectively) (Figs. 5C & D). The serial volumetric assessment for 20-day treatment revealed a marked attenuation of tumor progression by SH6 (Fig. 5E), with no obvious toxicity or weight loss being observed in the treated groups (Fig. EV5)

Next, we investigated the in vitro microsomal stability and pharmacokinetic (PK) properties of SH6. Incubation with human liver microsomes revealed a half-life of 32.4 min, compared to 7.3 min for the rapidly cleared positive control verapamil (Fig. 5F). To further study the suitability of SH6 as a drug candidate, we assessed the effect of SH6 in blocking hERG potassium channels, which could cause life-threatening arrhythmias(Sanguinetti & Tristani-Firouzi, 2006) (Fig. 5G). SH6 or the positive control amitriptyline were investigated at concentrations of 0.3, 1, 3,10, and 30 µM. It was found that Amitriptyline blocked hERG at an IC_50_ of 2.76μM, while SH6 blocked hERG at a much higher IC_50_ of 22.2 μM.

The *in vivo* Pharmacokinetic profile was next accessed to understand the bioavailability of SH6. Specifically, SH6 was administered to Swiss albino mice by intravenous injection (5mg/kg) or as an oral gavage (15mg/kg). Blood samples were collected at time intervals of 0.083, 0.25, 0.5, 1, 2, 4, 6, 8, and 24 hours for quantification of drug by LC-MS/MS (Fig. 5h). As seen from Fig. 5H, SH6 has an acceptable PK profile characterized by good oral bioavailability (63.6%) and a reasonable half-life of 1.67 hours. Importantly, SH6 was well tolerated in the treated mice and no mortality was observed at the doses employed.

## Discussion

The ability to degrade a protein of interest opens up a plethora of possibilities in designing degraders. The development of JQ1(Filippakopoulos et al., 2010), ER-PROTAC (Clinical trials NCT04072952), and AR-PROTAC (Clinical Trials NCT03888612), focuses attention of the drug development community to the advantages of harnessing the E3-ubiquitin-proteosome pathway as a therapeutic target(Henley & Koehler, 2021). This strategy has recently been employed to degrade transcription factors(Choi, Mahankali et al., 2017, Verhoeven et al., 2020). As a transcription factor, the uniqueness of SALL4 lies in the fact that it is expressed in embryonic cells, downregulated during development, absent in most adult tissues, and re-expressed in a significant fraction of almost all human adult tumors(Moein, Tenen et al., 2022, Tatetsu, Kong et al., 2016). The reappearance of SALL4 is associated with drug resistance in leukemia, lung, and breast cancer, and the onset of progenitor type hepatocellular carcinoma(Yong et al., 2013). The exclusive expression of SALL4 in cancer and not in adjacent untransformed cells provides an excellent therapeutic window that is rarely seen in drug targets. Immunomodulatory imide (IMiDs) drugs such as Thalidomide have been shown to degrade SALL4(Donovan et al., 2018, Matyskiela, Couto et al., 2018). It was reported that 5-hydroxythalidomide, a metabolite of Thalidomide, targets the ZFC2 of SALL4 to Cereblon(Furihata et al., 2020). These studies explained, at least in part, Thalidomide-induced birth defects, but failed to explain the lack of potency of IMiDs in SALL4 expressing cancer cell lines. We have observed that IMiDs only degrade SALL4 isoform A and not SALL4 isoform B. Here, we used the newly characterized SALL4 DNA binding crystal structure, a domain shared by SALL4A and B, the artificial intelligence program AlphaFold2, as well as a cell-based assay to screen for compounds that only target cancer cells that are SALL4 positive. This strategy also helps to reduce cytotoxicity issue by maximizing the chance of a high therapeutic index. Using this screening approach, we identified the non-IMiD lead compound SH6, which degrades the SALL4 protein and selectively suppresses the viability of SALL4 expressing cancer cells. The AlphaFold 3D structure prediction tool provides confidence scores for each residue based on the local distance difference test score. These models have been useful in drug discovery(Flower & Hurley, 2021), identifying pathogenic mutations(Bryant et al., 2022), and to investigate protein-protein interactions(Porta-Pardo et al., 2022). Using a combination of crystallography of the ZFC4 DNA binding regions with AlphaFold2, we predicted and demonstrated that SH6 binding to SALL4A and SALL4B differed from the previously reported Thalidomide binding site (i.e. cerebelon-SALL4A interface)(Furihata et al., 2020, Matyskiela et al., 2020), indicating that SH6 binds and functions differently from IMiDs.

In our study, we elected to employ the SALL4A and SALL4B AlphaFold models rather than the ZFC4:DNA crystal structure for the purposes of virtual screening, and this decision was predicated upon several critical considerations. While the ZFC4:DNA crystal structure shares a binding pocket region with the SALL4A/4B AlphaFold models, it is crucial to acknowledge that the ZFC4:DNA crystal structure only utilized a small segment of the protein. Specifically, it included only ZFC4, a mere 54 amino acids out of the 1053 amino acids in SALL4A or 616 amino acids in SALL4B. In contrast, the AlphaFold model utilizes the entire protein, encompassing regions that may exhibit various levels of structural flexibility, thereby affording a potentially more comprehensive representation of the molecular landscape for our virtual screening analysis. Furthermore, within the AlphaFold model, the binding site region is characterized by minimal or no exposure to water. This enhanced structural integrity within the binding site strengthens the predictive ability of our molecular docking investigations. In contrast, the ZFC4:DNA crystal structure exposes a substantial portion (25-30%) of the binding site to water. This incompleteness in the binding site representation could restrict the ability to fully explore the intricate interactions occurring within the binding site. Exposure of a protein’s binding pocket to water typically diminishes ligand binding affinity due to the hydrophobic effect. Hydrophobic regions within the pocket tend to interact favourably with the ligand’s hydrophobic moieties. Water presence energetically hinders ligand binding, a crucial consideration in molecular docking studies. In summary, AlphaFold models provided a more comprehensive representation of the protein’s structure and binding site, making them well-suited for docking and virtual screening applications.

Cell-based screening was then employed to evaluate the selected compounds in targeting SALL4 expressing cancer cells. To ensure that the cell lines selected are representative and relevant for SALL4 study, we used cell lines with high and low endogenous SALL4 expression, as well isogenic cell lines overexpressing SALL4A or SALL4B (SNU387-TgSALL4A and SNU387-TgSALL4B)(Tan et al., 2019). This five-cell line cell-based screening platform has led to the discovery of the role of SALL4 in inhibition of oxidative phosphorylation and opens a new potential strategy to target SALL4 positive cancers(Tan et al., 2019, Yong et al., 2013). In this approach, SH6 showed selectivity against SNU398 cells when compared to SNU387 cells. Furthermore, it was found that SNU387-TgSALL4A and SNU387-TgSALL4B were highly sensitive to SH6 treatment compared to SNU387 wild type isogenic line. Overall, our phenotypic screening results support that SH6 can selectively target cancer cell survival mediated by SALL4.

More recently, in other studies, the dependency on SALL4 in SNU398 cells was demonstrated through a shRNA-medicated knock down approach(Vu et al., 2023) where we down-regulated either total SALL4 (both A and B) or SALL4B only in SNU398 cells. We observed that upon SALL4/SALL4B-specific knockdown, the apoptotic population of SNU398 cells was increased by 40%, and the cellular growth ability of these cells were significantly inhibited tested by soft agar colony formation and clonogenic assays. In comparison, down-regulation of SALL4 had no effect on SNU387 cells.

An unforeseen discovery was made when we observed that SALL4 degradation caused by SH6 could be reversed with the use of proteasomal inhibitors MG132 and MLN4924. Despite the structural differences between SH6 and IMiDs, we did not anticipate the involvement of a similar E3 degradation pathway. Cullin based E3 ligases need to be neddylated by NEDD8 in order to activate their holoenzyme ubiquitin ligase activities(Soucy, Smith et al., 2009). The neddylation itself is catalysed by E1 ligase NAE, in which the formation of a NAE-NEDD8 thioester is crucial for subsequent transfer of NEDD8 to E2s and E3s. MLN4924 forms a covalent adduct with NEDD8 and inactivates it, blocking the very first step of NAE-NEDD8 formation. When SALL4 high cells are treated with MLN4924, SALL4 protein expression is rescued. Furthermore, viability of SALL4 high cells, which is severely impaired by SH6, is rescued when co-treated with MLN4924 (Figs. 3E). These results indicate that SH6 mediated SALL4 degradation involves the cullin-based E3 system. Moreover, when we employed in-cell CETSA to investigate cell-based target engagement by SH6, we found that SH6 could engage SALL4, CUL4A, and CRBN, causing significant higher thermal stability for these proteins (Fig. 3G). A potential model has been proposed (see Graphical Abstract), in which IMiDs target ZFC2 of SALL4A, which is not present in SALL4B. This results in the inability of IMiDs to target SALL4+ cells which express both isoforms. In contrast, SH6 targets ZFC4, which is present in both SALL4A and SALL4B isoforms, and directs it to the CUL4-RBX1-DDB1-CRBN-ubiquitination based degradation pathway, resulting in death of SALL4+ cancer cells.

Importantly, the degradation of SALL4 protein requires ZFC4 (Fig. 4A). Mass spectrometry analysis of the SH6 degrome searching for additional ZFC4 targets demonstrated that among 294 down-regulated proteins, 2 proteins were identified to have ZFC4 sequences, ZBTB7A and ZBTB11 (Figs. 4B,C & D). Both proteins were confirmed to be down-regulated by treatment with SH6 but not with the IMiD Pomalidomide (Fig. 4E). ZBTB7A has a crystal structure available for comparison to SALL4 ZFC4, and we found that the two crystals match well (Fig. 4F). Overall, our findings support that SH6 is an IMiD-independent ZFC4 degrader.

Herein, we developed a small molecule that degrades both SALL4 isoforms and showed prominent and specific efficacy in SALL4 expressing cancer cells. Furthermore, our xenograft studies demonstrate that SH6 inhibits tumour growth without causing toxicity to the mice. Cell line studies also indicate that SH6 is selective against SALL4 expressing and not SALL4 negative cancer cells or to untransformed liver cells. Unlike traditional PROTACs that are bulky, SH6 presents a desirable pharmacokinetic profile and oral bioavailability, and shows exceptional antitumor effects in murine xenograft studies.

In summary, our study showcases the feasibility of combining crystallographic structural information of a zinc finger DNA binding transcription factor and AlphaFold2(Jumper et al., 2021, Jumper & Hassabis, 2022), along with a phenotypic screen, to discover a targeted therapeutic molecule against the previously un-targetable oncofetal protein SALL4. We anticipate future medicinal chemistry studies to develop SH6 from a research tool compound to that of a clinically therapeutic drug targeting cancers with SALL4 expression. In addition, the discovery of the ZFC4 binding site has expanded the screening potential of existing and novel compound libraries, enabling the identification of compounds that can target proteins containing ZFC4, a new degron that not covered by IMiDs.

## Materials and Methods

### Protein expression and purification

Residues 864-929 of human SALL4 were inserted into the pGEX4T1 (N-terminal GST thrombin tag) vector. A N-terminal GST tagged construct of human SALL4 including residues 864-929 (RRQA**<u>KQHGCTRCGKNFSSASALQIHERTHTGEKPFVCNICGRAFTTKGNLKVHYMTHG</u>**ANNNSARR, with SALL4 864-929 denoted in bold and underlined font) was overexpressed in E. coli BL21 (DE3) and purified using affinity chromatography and size-exclusion chromatography. Briefly, cells were grown at 37°C in TB medium in the presence of 50 μg/ml of ampicillin to an OD of 0.8, cooled to 17°C, induced with 500 μM isopropyl-1-thio-D-galactopyranoside (IPTG), incubated overnight at 17°C, collected by centrifugation, and stored at -80°C. Cell pellets were lysed in buffer A (25 mM HEPES, pH 7.5, 200 mM NaCl, 7 mM mercapto-ethanol, and 10 μM zinc chloride) using Microfluidizer (Microfluidics), and the resulting lysate was centrifuged at 30,000g for 40 min. Glutathione superflow agarose beads (Fisher) were mixed with cleared lysate for 90 min and washed with buffer A. Beads were transferred to an FPLC-compatible column, and the bound protein was washed further with buffer A for 10 column volumes and eluted with buffer B (25 mM HEPES, pH 7.5, 200 mM NaCl, 7 mM mercapto-ethanol, 10 μM zinc chloride, and 15 mM glutathione). To cleave the GST-tag, thrombin was added to the eluted sample and incubated overnight at 4°C. The resulting solution was concentrated and purified further using a Superdex 75 16/600 column (Cytiva) in buffer C containing 20 mM HEPES, pH 7.5, 200 mM NaCl, 0.5 mM TCEP, 1 mM DTT, and 10 μM zinc chloride. SALL4 containing fractions were pooled, concentrated to ∼6mg/mL, and stored in -80°C.

### Crystallization

Using Formulatrix NT8 and ArtRobbins Phoenix liquid handlers, we pre-incubated 100 nl samples of 500uM SALL4 with 750uM 12-base pair blunt-end duplex DNA (CGAAATATTAGC) for 1 hour. Subsequently, these samples were dispensed in an equal volume of crystallization buffer (comprising 30% PEG3350, 0.04M NH4SO4, and BisTris at pH 6.0) and incubated against 25 ul of reservoir crystallization buffer in a 384-well hanging-drop vapor diffusion microtiter plate. The samples were incubated for three days at 20°C and observed using Formulatrix Rock Imager. The crystals were briefly transferred into a crystallization buffer enriched with 25% glycerol before being flash-frozen in liquid nitrogen. They were then shipped to the synchrotron facility for data collection.

### Data collection and structure determination

Diffraction data were collected at beamline 24ID-E of the NE-CAT at the Advanced Photon Source (Argonne National Laboratory). Data sets were integrated and scaled using XDS(Kabsch, 2010). Structures were solved by SAD-phasing using the program SOLVE(Adams, Afonine et al., 2010) Iterative manual model building and refinement using Phenix(Adams et al., 2010) and Coot(Emsley & Cowtan, 2004)led to a model with excellent statistics. Crystal structures, with statistics, shown in Table EV1, were deposited into the Protein Data Bank (PDB code 8CUC) with the PDB validation report.

### AlphaFold2

The 3D structural models of SALL4A and SALL4B were generated using AlphaFold2(Jumper et al., 2021, Jumper & Hassabis, 2022) through Colab python notebook(Matyskiela et al., 2018). Starting from the sequences of SALL4A and SALL4B, AlphaFold2 was run using “none” or “pdb70” template mode, MSA mode “mmseqs2 (Uniref + environmental), and pair mode “paired+unpaired” options. AlphaFold generated five structural models. The best ranked model by average pLDDT score was used for further analysis.

### Molecular docking prediction

Since AlphaFold models do not include ligands, SiteMap(Schrödinger, 2020d) (https://colab.research.google.com/github/sokrypton/ColabFold/blob/main/AlphaFold2.ipynb) was used to predict potential binding sites (Schrödinger 2020). To visualize and evaluate top binding sites, probable binding site regions with a minimum of fifteen site points were generated in a more restrictive hydrophobic environment using a standard grid. Based on Dscore and druggability score calculated by SiteMap, four pockets are considered for SALL4A and one pocket for SALL4B for further molecular docking analysis.

The AlphaFold models and crystal structure were prepared by a standard protocol protein preparation wizard module(Schrödinger, 2020c), which adds hydrogen atoms, repairs imperfect side chains and assigns protonated states of the system at pH 7.0 ± 2.0. The receptor grids for all proteins were generated on predicted binding sites from site map analysis using the GLIDE module(Schrödinger, 2020a) of Schrodinger. The generated site map points were used to generate a grid box 10 Å in size. Molecular docking was carried out using GLIDE module. Before docking, The LigPrep module(Schrödinger, 2020b) was utilized to prepare the ligands at pH = 7.5 ± 1, for tautomer generation, and energy minimization with the OPLS3e force field. Ligands were docked on predicted binding sites in the standard precision (SP) mode to generate 10 poses per ligand through flexible ligand sampling. Their binding poses were evaluated using Glide SP scoring functions.

The ZFC2-4 of SALL4A and ZFC4 of SALL4B models were docked to CUL4A using Piper(Kozakov, Brenke et al., 2006). 70000 rotations were sampled, and the top 30 poses were returned from each docking job. We considered the best scored poses for further analysis.

### Cell based phenotypic screen

Empty vector, SALL4A, and SALL4B expressing isogenic cell lines were generated by transducing the SALL4 negative hepatocellular carcinoma SNU-387 cells with empty vector, SALL4A, or SALL4B FUW-Luc-mCh-puro lentiviral constructs. Cells were plated in 50 μl of RPMI culture media in 384-well white flat-bottom plates (Corning) and incubated at 37°C in a humidified atmosphere of 5% CO2 overnight. Cell numbers per well were 1500 for SNU-398, and 750 for SNU-387 and SNU-387 isogenic lines. After overnight incubation, varying concentrations of compounds 1-80 were added to cells with multichannel electronic pipettes (Rainin). Cells were then incubated for 72 hrs at 37°C in a humidified atmosphere of 5% CO2 before 10 μl of CellTiter-Glo reagent was added to the wells with the MultiFlo Microplate Dispenser (BioTek). Cells were incubated at room temperature for a minimum of 10 minutes after which luminescence readings were recorded by an Infinite M1000 Microplate Reader (Tecan). Each data point was performed in n=3 independent cell cultures. IC50 of compounds were determined using nonlinear regression fitting from Prism (GraphPad).

### Western blots

SNU398 high SALL4 HCC cells were incubated with the compounds (SH6 or pomalidomide) for concentration and times indicated in the figure legends. Cells were then harvested by cell scraper and washed with PBS. The collected pellets were lysed with RIPA buffer (50 mM Tris, 150 mM NaCl, 1% TritonX-100, 0.5% sodium deoxycholate, and 0.1% SDS) supplemented with Complete^TM^ protease inhibitor cocktail (Roche, Switzerland). The extracted protein lysates were denatured with 4X SDS sample buffer (200mM Tris-HCl pH 6.8, 8% SDS, 40% glycerol, 4% β-mercaptoethanol, 50mM EDTA, 0.08% bromophenol blue) at 99 °C for 5 minutes. Equal amounts of protein were subjected to electrophoresis in 8% SDS-PAGE gels, and then transferred to PVDF membranes. After blocking in Blocking One (Nacalai Tesque), the membrane was probed with primary antibodies to SALL4 (Santa Cruz Biotechnology, sc-101147), β-Actin (Santa Cruz Biotechnology, sc-47778), Cul4A (#2699) Cell Signalling Technology, CRBN (D8H3S, #71810) Cell Signalling Technology, LRF (13E9, 14-3309) from eBioscience, Armenian hamster IgG-HRP (sc-2789) from Santa Cruz, and ZBTB11 antibody (25215-1-AP) from Proteintech, and incubated overnight at 4 °C.

After washing with TBS-T, membranes were incubated with secondary HRP-conjugated antibody to murine (Santa Cruz Biotechnology, sc-2005), or Armenian hamster IgG-HRP (Santa Cruz Biotechnology, sc-2789) for 1 hour. Luminata™ western HRP substrate (Millipore) was applied to the membrane for visualization.

### Dual-Luciferase Reporter Assay

SALL4B was subcloned into Nluc/Fluc plasmid(pLL3.7-EF1a-IRES-Gateway-nluc-2xHA-IRES2-fluc-hCL1-P2A-Puro) provided by William Kaelin(Lu et al., 2014) and stably transfected into H1299 cells (H1299-SALL4BNLuc). H1299-SALL4BNLuc cells were seeded at 1000 cells per well in 384 well plates, and treated with drugs for 24hrs before dual luciferase assays were performed using the Nano-Glo®Dual-Luciferase® Reporter Assay System (Promega, Cat#: N1650) as described by the manufacturer. Luminescence was read by Envision^TM^ (Perkin Elmer) after 24 hours of drug treatment. Data was presented by comparing DMSO normalized Nanoluc luminescence (NLuc) to DMSO normalized Firefly luminescence (FLuc). Biological triplicates were performed for each drug concentration. Each data point was performed in n=3 independent cell cultures. IC50 of compounds were determined using nonlinear regression fitting from Prism (GraphPad).

### Cell Culture

All HCC cell lines were obtained from ATCC and grown according to the provider’s instructions in the absence of antibiotics. Human hepatocellular carcinoma cell lines (SNU398, SNU387, isogeneic lines made as described above, A549, H1299, and H661) were maintained in Dulbecco’s Modified Eagle’s medium (DMEM) and Roswell Park Memorial Institute 1640 medium (RPMI) (Life Technologies, Carlsbad, CA) with 10% fetal bovine serum (FBS) (Invitrogen) and 2 mM L-Glutamine (Invitrogen). These cell lines were cultured at 37°C in a humidified incubator with 5% CO2.

### Cellular Thermal Shift Assay (CETSA)

Cells were treated with 50µM SH6 or DMSO for 4 hours in 37°C with 5% of CO_2_. Cells were harvested and lysed in 1X NP-40 lysis buffer containing 10mM Tris-HCl (pH 7.4), 150mM NaCl, 2.7mM KCl and 0.4% NP-40. The mixture was then rotated end-over-end for 30 minutes at 4°C for lysis. The lysate was separated from the cell debris by centrifugation at 20,000 rpm for 10mins at 4°C. Lysates were heated in a thermal cycler at different temperatures, ranging from 45°C to 57°C for 3 minutes, followed by a 3-minute cool-down at room temperature. The heated lysates were centrifuged at 15,000 rpm for 10 minutes at 4°C to separate the precipitates from the soluble fractions. The supernatants were subjected to western blot analysis with CUL4A (Cell Signalling Technology-2699), NEDD8 (ThermoFisher Scientific-PA517476), SALL4 (Santa Cruz Biotechnology, sc-101147), and CRBN (Cell Signalling Technology-71810).

### Mass Spec Analysis

In brief, SNU398 was grown treated with SH6 (50uM) or DMSO for 24hr. Samples were extracted in quadruplicates using RIPA buffer (Sigma) according to the manufacturer’s instructions. In total, 200 μg of protein was used for MS sample preparations as previously described 1. Samples were boiled at 95 °C prior to separation on a 12% NuPAGE Bis-Tris precast gel (Thermo Fisher Scientific) for 15 min at 170 V in 1× MOPS buffer. The gel was fixed using the Colloidal Blue Staining Kit (Thermo Fisher Scientific), and each lane was divided into 2 equal fractions. For in-gel digestion, samples were destained in destaining buffer (25 mM ammonium bicarbonate; 50% ethanol), reduced in 10 mM DTT for 1 h at 56 °C followed by alkylation with 55 mM iodoacetamide (Sigma) for 45 min in the dark. Tryptic digest was performed in 50 mM ammonium bicarbonate buffer with 2 μg trypsin (Promega) at 37 °C overnight. Peptides were desalted on StageTips and analyzed by nanoflow liquid chromatography on an EASY-nLC 1200 system coupled to a Q Exactive HF mass spectrometer (Thermo Fisher Scientific). Peptides were separated on a C18-reversed phase column (25 cm long, 75 μm inner diameter) packed in-house with ReproSil-Pur C18-AQ 1.9 μm resin (Dr Maisch). The column was mounted on an Easy Flex Nano Source and temperature controlled by a column oven (Sonation) at 40 °C. A 215-min gradient from 2 to 40% acetonitrile in 0.5% formic acid at a flow of 225 nl/min was used. Spray voltage was set to 2.4 kV. The Q Exactive HF was operated with a TOP20 MS/MS spectra acquisition method per MS full scan. MS scans were conducted with 60,000 at a maximum injection time of 20 ms and MS/MS scans with 15,000 resolution at a maximum injection time of 50 ms. The raw files were processed with MaxQuant version 1.5.2.8 using the LFQ quantification option2 on unique+razor peptides with at least 2 ratio counts. Carbamidomethylation was set as fixed modification while methionine oxidation and protein N-acetylation were considered as variable modifications. Search results were filtered with a false discovery rate of 0.01. Known contaminants, proteins groups only identified by site, and reverse hits of the MaxQuant results were removed. LFQ intensities can be found in Supplementary Table 4. Differential analysis (log base2 fold change < 2, adjusted p value < 0.05) was performed and downregulated protein were selected based on these criteria. Protein sequence of the down regulated proteins were compared and aligned to that of SALL4 ZFC4 to search for other ZFC4 targets of SH6.

### HCC Mouse Xenografts

Animals were maintained, and all animal work was performed following the protocols approved by the Children’s Hospital Boston Institutional Animal Care and Use Committee. The SNU-398 cell line was cultured as described above in ‘Cell Culture’. To establish the SNU-398 xenografts, NOD.Cg-Prkdc^scid^ Il2rg^tm1Wjl^/SzJ (NSG) female mice (6-8 weeks) were anesthetized using 2% isoflurane USP (Baxter), and 2 x 10^6^ SNU-398 cells in 200 ul of 1:1 RPMI:Matrigel were subcutaneously injected to the mouse flanks. When tumors were palpable at day 5, mice were randomized to receive either SH6 treatment or vehicle containing 5% DMSO in PBS. Freshly prepared vehicle or SH6 was administered by intraperitoneal (i.p.) injection, once daily (qd) at 1 and 2.5 mg/kg. Before each drug injection, mouse weight and tumor volume were monitored. Mice were euthanized once tumors reached more than 1.5 cm in diameter. The tumors were harvested, imaged, and snap-frozen for storage at -80°C until later use. All statistical analysis was performed using a one-way ANOVA and Student’s T-test. Error bars represents standard deviation of mean. Xenograft studies were performed twice independently. Representative data from a single experiment are shown.

### Synthesis of SH derivatives

Schematic:

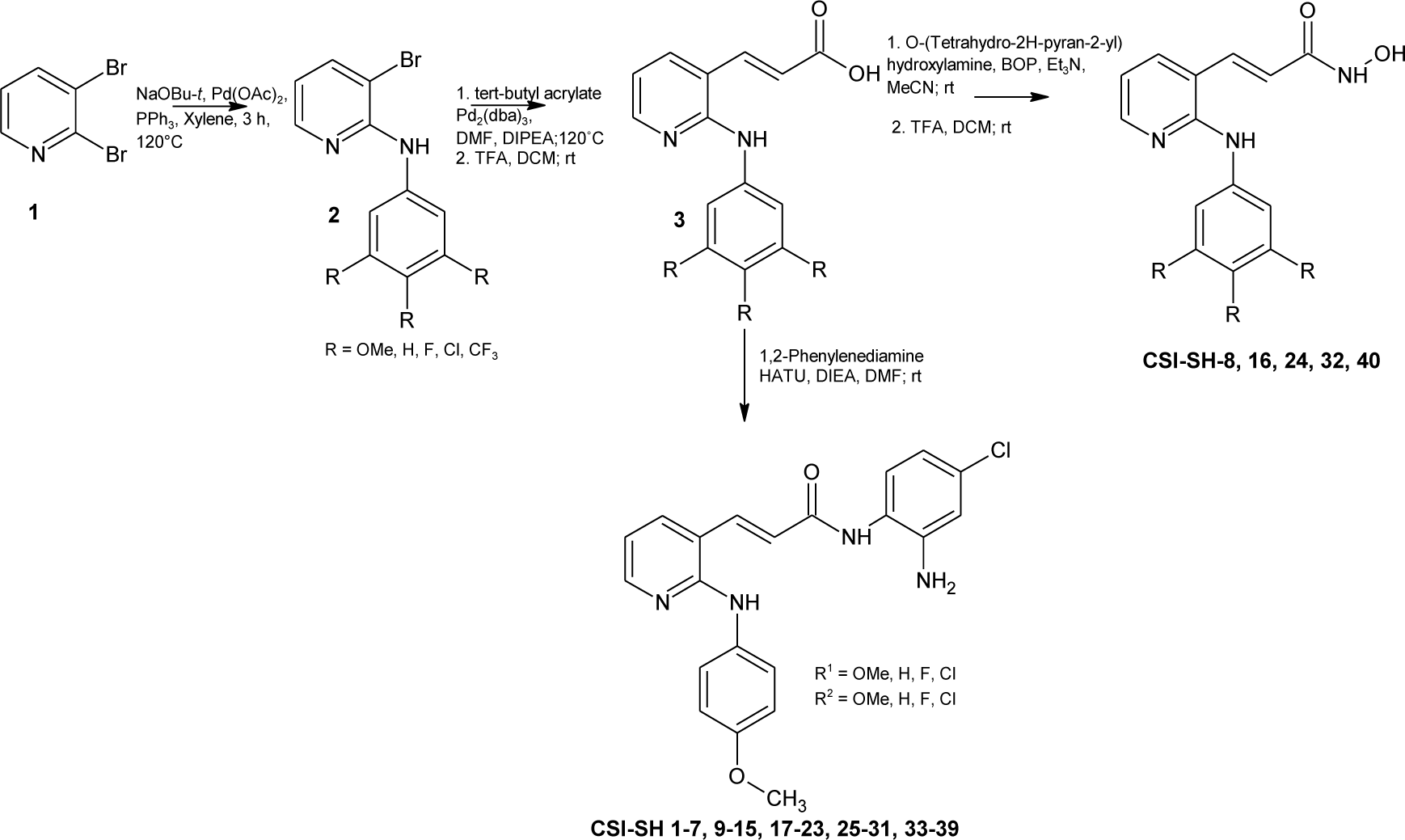

Synthesis of 2, 3-Bromo-N-(3,4,5-trimethoxypheny1)pyridine-2-amine: In a 5-ml screw-capped vial, palladium acetate (0.14 mg, MW-225, 0.62 mmol), 4,5-bis(diphenylphosphino)-9,9-dimethylxanthene (Xantphos) (0.73 g, MW-579, 1.26 mmol), cesium carbonate (8.25 g, MW-326, 25.2 mmol), 2,3-dibromopyridine (3.0 g, MW-237, 12.6 mmol) and degassed toluene (30 ml) were mixed, and to this, trimethoxyaniline (2.31 g, MW-183, 12.6 mmol) was added. Then, nitrogen gas was enclosed in the vial, and the vial was stoppered tightly, and the mixture was stirred with heating at an external temperature of 115°C for 1.5 hours. After completion of the reaction, the reaction solution was cooled to room temperature, toluene (150 ml) and water (150 ml) were added thereto, and the insoluble was filtered off through Celite. The filtrate was washed with toluene (300 ml), and then the obtained organic layer was washed with saturated brine (250 ml), dried over anhydrous sodium sulfate and concentrated under reduced pressure. To the residue, a mixture (500 ml) of hexane/ethyl acetate (5/1) was added. The mixture was stirred at room temperature for 10 minutes, and then the insoluble was filtered off. The filtrate was concentrated under reduced pressure, and dried under reduced pressure at 50°C, to yield the title compound (3.4 g, MW-339) as a pale yellow solid (yield 80%). ^1^H-NMR (CDCl_3_, 400 MHz) δ (ppm): 8.17(dd, 1H, *J*= 8.0, 4.0 Hz, 1H), 7.75 (dd, 1H, *J*= 8.0, 4.0 Hz), 6.91 (brs, 1H), 6.91 (s, 2H), 6.65 (m, 1H), 3.89 (s, 6H), 3.83 (s, 3H). ^13^C-NMR (CDCl_3_, 400 MHz) δ (ppm): 153.35, 152.02, 146.49, 140.25, 135.72, 134.05, 115.51, 106.30, 98.34, 60.94, 56.13.

Synthesis of 3, (2E)-3-[2-(3,4,5-trimethoxyanilino)pyridin-3-yl]prop-2-enoic acid *t*-butyl ester: In a 100 mL flask, a mixture of 2 (880mg, 5mmol), *tert*-butylacrylate (4 mL, 27.5 mmol), Diisopropyl ethylamine (4 mL, 23 mmol), tri-*o*-tolylphosphine (0.95 g, 3 mmol), Pd_2_(dba)_3_ (0.375 g, 0.4 mmol) in anhydrous DMF (20 mL) was stirred at 120°C (preheated oil bath) for 2 h under nitrogen. After removal of DMF, the crude residue was purified using column chromatography (0 to 1 % DCM:MeOH) to afford 0.95 g of product. ^1^H-NMR (CDCl_3_, 400 MHz) δ (ppm): 8.25 (dd, 1H, *J*= 8.0, 4.0 Hz), 7.70 (m, 2H), 6.82 (m, 3H), 6.43 (s, 1H), 6.37 (d, 1H, *J*= 12 Hz), 3.86 (s, 6H), 3.83 (s, 3H), 1.54 (s, 9H). ^13^C-NMR (CDCl_3_, 400 MHz) δ (ppm): 165.85, 153.36, 153.31, 149.34, 137.48, 136.19, 136.17, 133.98, 123.26, 116.85, 115.52, 98.54, 81.05, 60.91, 56.11, 28.16.

Synthesis of 4, (2E)-3-[2-(3,4,5-trimethoxyanilino)pyridin-3-yl]prop-2-enoic acid: Ester 3 (0.9 g, mmol) dissolved in 40% TFA in dichloromethane (50 mL) and the solution was stirred at room temperature overnight. The solvent was removed under vacuum with acetonitrile (3x30 mL) and stored under high vacuum for 12h. The solid residue 4 (0.72 g, 93%) was used for coupling with amines as such without further purification. ^1^H-NMR (CDCl_3_, 400 MHz) δ (ppm): 9.03(brs, 1H), 8.09 (m, 2H), 7.91 (d, 1H, *J*= 8.0 Hz), 6.88 (brs, 3H), 6.50 (m, 1H), 3.75 (s, 6H), 3.66 (s, 3H).

Synthesis of CSI-SH-6, (2E)-N-(2-amino-4-chlorophenyl)-3-[2-(3, 4, 5-trimethoxy anilino) pyridin-3-yl] prop-2-enamide: In a 100 mL RB flask acid 4 (MW-330, 1g, 3.03 mmol) was added DMF (15 mL), HATU (MW-380, 1.4 g, 3.6 mmol), diisopropyl ethylamine (129.25, 2.10 ml, 12.5 mmol) and stirred for 15 min. Then the corresponding amine, chlorophenylenediamine (MW-142, 0.4g, 2.83 mmol) was added to the reaction mixture and the solution was stirred at 22 °C for 21 h. Then H_2_O (20 mL) was added; the resulting precipitate was isolated by vacuum filtration and purified by silica gel chromatography (97:3:: CH_2_Cl_2_/ MeOH) to afford the desired product as a pale yellow solid. Mol Wt-454. ^1^H-NMR (DMSO-D_6_, 400 MHz) δ (ppm): 9.41 (brs, 1H), 8.54 (s, 1H), 8.19 (d, 1H, *J* = 4 Hz), 7.87 (m, 2H), 7.41 (d, 1H, *J* = 8 Hz), 7.00 (s, 2H), 6.84 (m, 3 Hz), 6.61 (dd, 1H, *J* = 8, 2.4 Hz), 5.30 (brs, 2H), 3.74 (s, 6H), 3.62 (s, 3H).

## Acknowledgements

This work was supported by the Singapore Ministry of Health’s National Medical Research Council (Singapore Translational Research (STaR) Investigator Award STaR18nov-0002 (D.G.T.)); the Singapore Ministry of Education under its Research Centres of Excellence initiative; NIH/NHLBI P01HL131477-01A1 (D.G.T); the Xiu research fund and AGA/Jenzabar research fund (L.C.); the Northeastern Collaborative Access Team beamlines, funded by the National Institute of General Medical Sciences from the National Institutes of Health (P30 GM124165); the Eiger 16M detector on the 24-ID-E beam line funded by a NIH-ORIP HEI grant (S10OD021527); and resources of the Advanced Photon Source, a U.S. Department of Energy (DOE) Office of Science User Facility operated for the DOE Office of Science by Argonne National Laboratory under Contract No. DE-AC02-06CH11357. S.D.P acknowledges funding from from the Linde Family Foundation, NIBR, 3DC, and DDCF. We thank William Kaelin for the Nluc/Fluc plasmid.

## Author contributions

Conceptualization: BHL, CL, DGT

Methodology: BHL, SR, CKJ, CG, HF, JPT, KK, KVL, HSS, KS, XT, LF, JT, HA, JQ, SDP

Investigation: BHL, SR, CKJ, CG, HF, JPT, KK, KVL, HSS, KS, XT, LF, JT, MB, HA, JQ, SDP

Supervision: BHL, SR, CL, DGT, ML, MLG, HF, HA, SDP

Writing—original draft: BHL, SR, CL, DGT

Writing—review & editing: BHL, SR, CL, DGT, ML, CKJ, CG, MLG, HF

## Code availability

No unique code was developed for this study.

## Data availability

Crystal structure statistic was deposited into the Protein Data Bank (PDB code 8CUC).

---

**Figure EV1.**
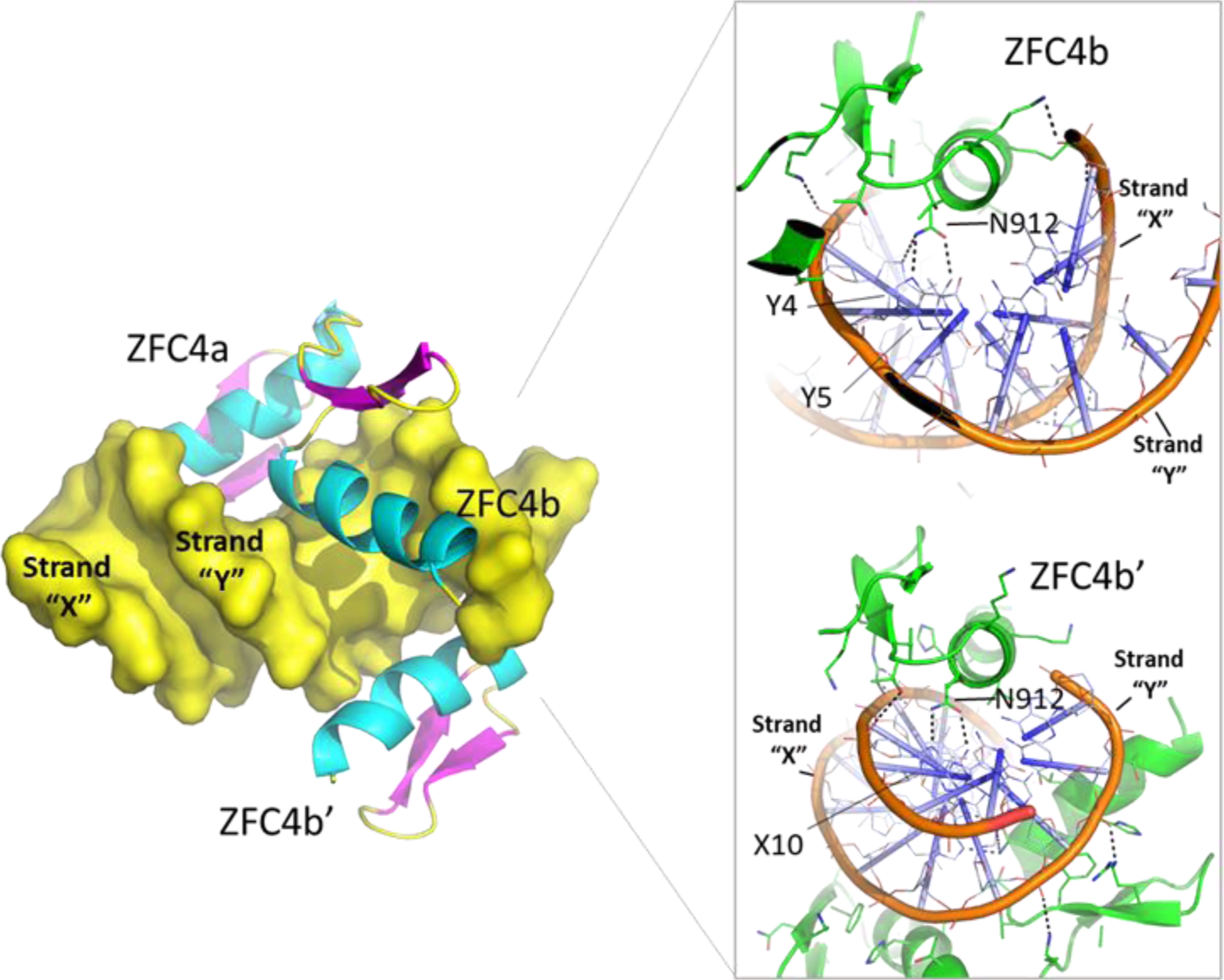
Crystal Structure of the fourth SALL4 zinc finger cluster (ZFC4, teal and magenta) bound to DNA (yellow). The structural view demonstrates how the two zinc fingers of the first SALL4 molecule (ZFC4b & ZFC4b’) and the second zinc finger of the second SALL4 molecule (ZFC4a) in the asymmetric unit bind in the major groove of DNA. Structural view in the box depicting the specific interactions between N912 of SALL4 and adenines at position Y4, Y5 and X10 of DNA.

**Figure EV2.**
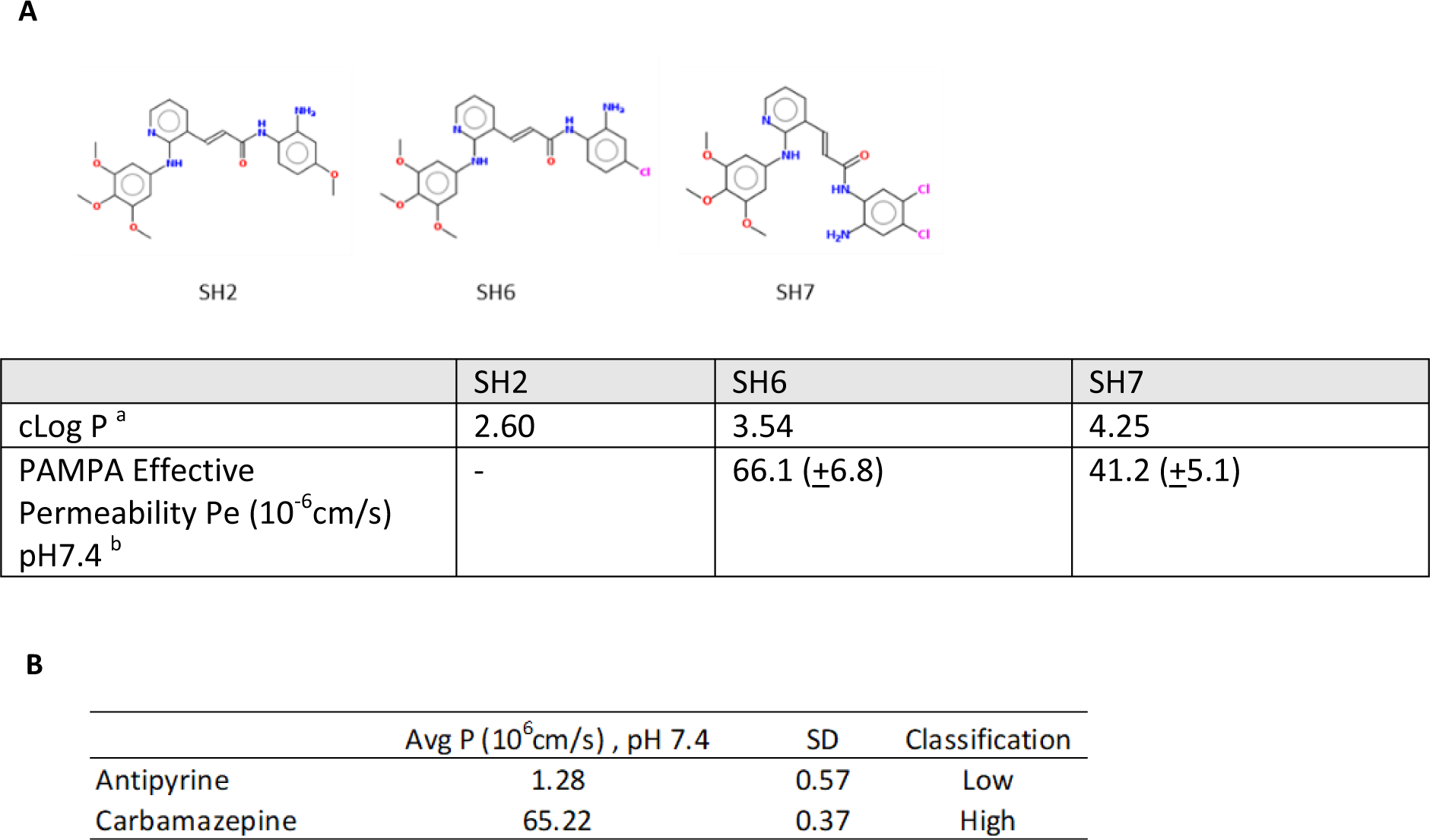
Structures and physicochemical properties of potent analogues SH2, SH6, and SH7. a) These compounds have the lowest EC50 values (< 3 uM) on SALL4 high SNU 398 cells b) Permeability test (PAMPA) result of negative control (Antipyrine), and positive control (Carbamazepine).

**Figure EV3.**
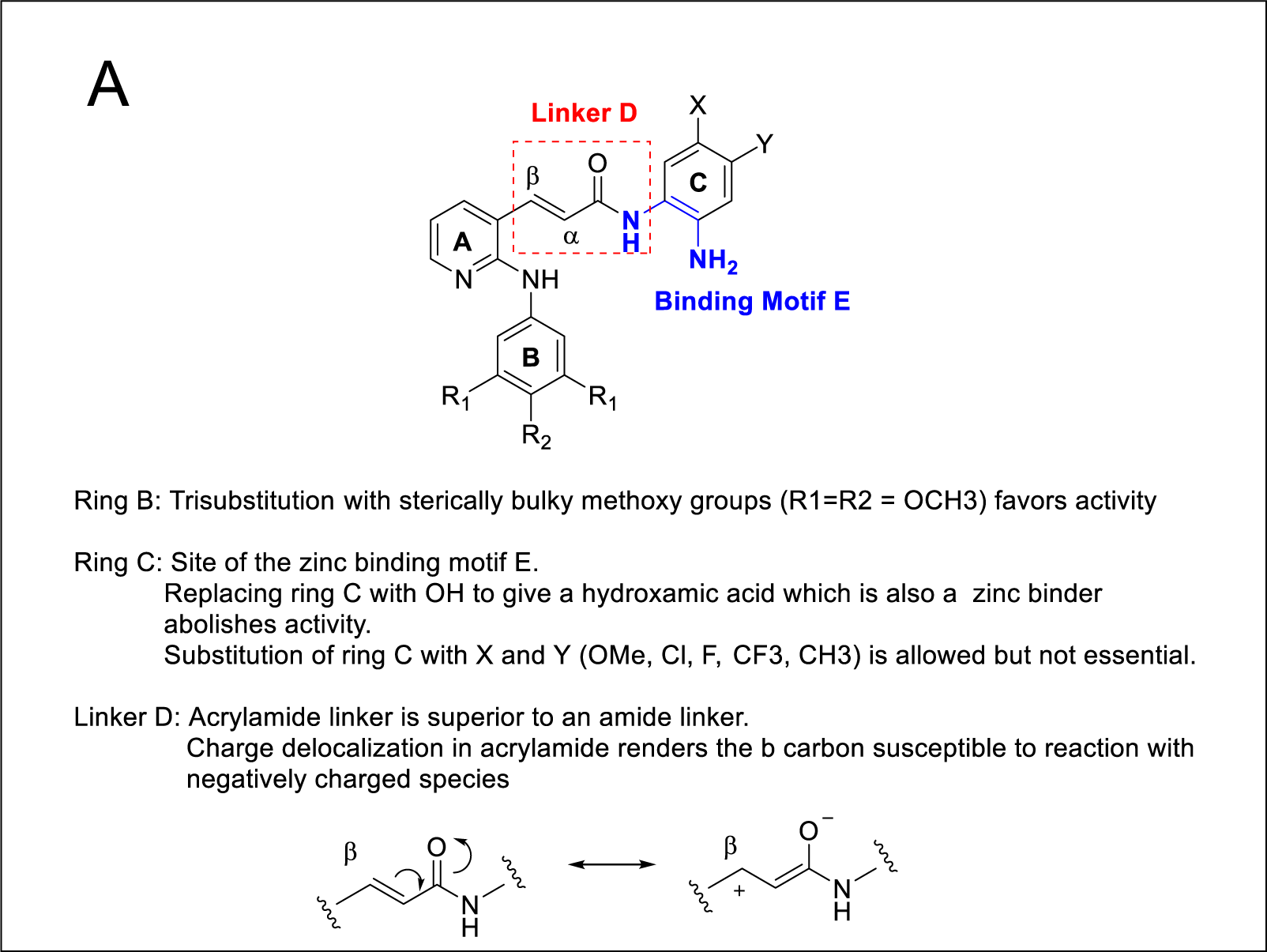
Structural Activity of SH library. Three key structure-activity correlations were noted. First, the acrylamide linker is an indispensable feature for activity. Replacing this linker with an amide resulted in SH40-SH47, SH58 and SH 74 which have markedly lower activity against the SALL4 high cells. In fact, of the 40 SH compounds (SH41-SH80) with amide linkers, only three have EC50 values ≤20 uM on SALL4 high cells, as compared to 31 out of 40 compounds (SH1-SH40) bearing the acrylamide linker. Unlike the amide, extensive delocalization of the negative charge occurs in the acrylamide moiety, resulting in an electron deficient β carbon which would in turn be more susceptible to reaction with electron rich species. It is conceivable that this enhanced reactivity contributed in some way to the greater cell-based potencies of the acrylamide based SH compounds. A second requirement for potent activity is the phenylene diamine ring D in the HDAC component of the hybrid design. The two amino groups on ring D are ortho to each other, with one amino embedded within the acrylamide linker while the other is a free amine. The ortho positioning of the amino groups is optimal for zinc binding. When Ring D is replaced by hydroxyl (OH) as seen in SH8, SH16, SH24, SH32 and SH40, activity was completely abolished, notwithstanding the zinc binding capability of the replacement N-hydroxyacrylamide moiety and the presence of the acrylamide linker. The diminished activity is likely due to the absence of steric bulk in the replacement moiety. The phenylene diamine Ring D is substantially larger in size and its steric bulk is further augmented by mono- or di-substitution with halogens (chloro, fluoro), methyl or methoxy groups. However, the contribution of these substituents to activity is not comparable to that of the acrylamide linker as it is noted that the absence of substituents on Ring D as seen in SH1, did not abolish its activity (EC50 of 3 uM). Lastly, we noted that 7 of the 9 potent compounds (EC50 < 3 uM) have a trimethoxy substituted ring B. Rings A and B are part of the scaffold found in the antimicrotubule agent E7010, with the anilino ring B attached to the ortho position of pyridyl ring A. Mono-substitution of ring B with either methoxy, chloro, fluoro or trifluoromethyl generally led to significant losses in activity, with the exception of SH18 and SH34. It may be that the sterically bulky and lipophilic trimethoxy substituted ring B is required to establish strong interactions (van der Waals, hydrophobic) with a putative receptor, hence contributing to enhanced potency.

**Figure EV4.**
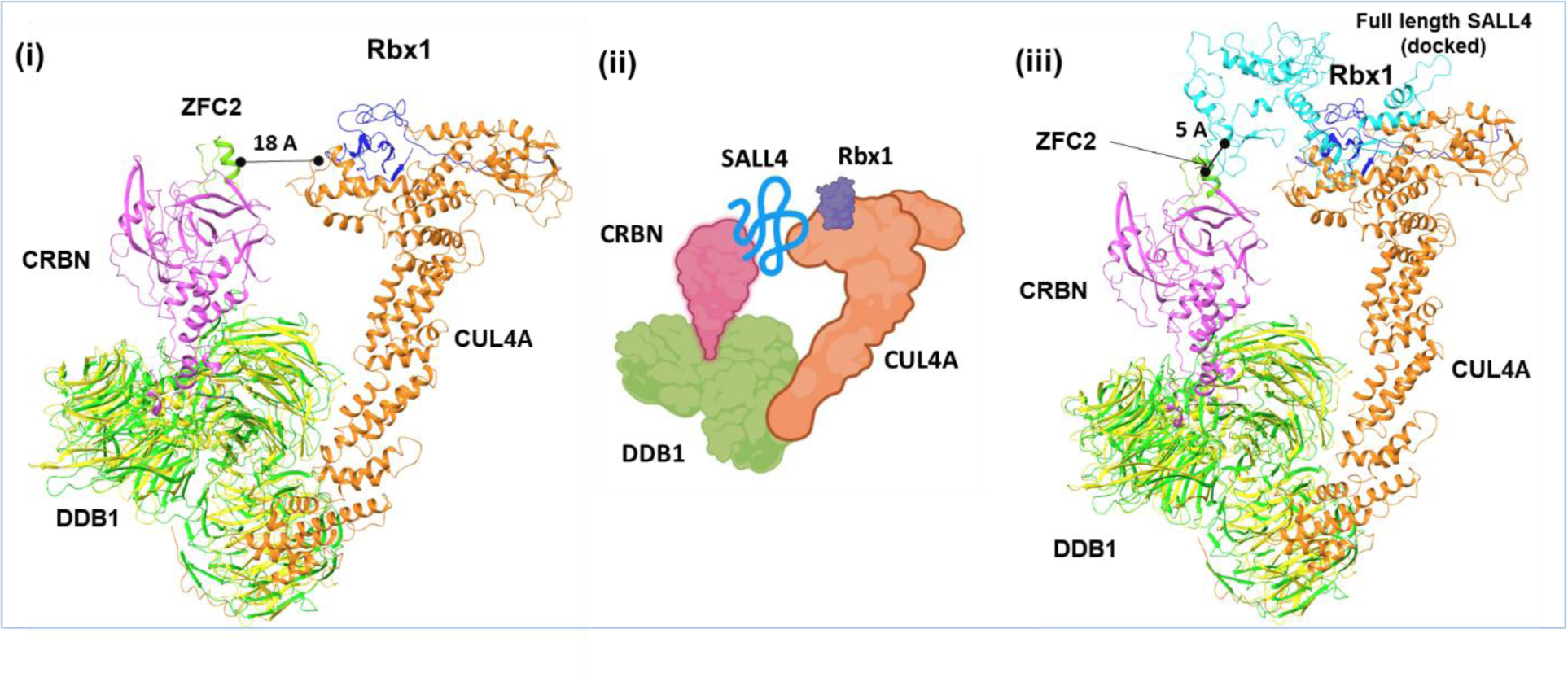
The structural model of SALL4A recruited to CRBN-DDB1-CUL4A-Rbx1 complex. ZFC2-4 was predicted to bind to CUL4A near the Rbx1 site. (A) Superimposed 3D structures of the SALL4A ZnF3-CRBN-DDB1(green) complex (PDB ID: 6UML) and DDB1(yellow)-CUL4A-Rbx1 complex (PDB ID: 2HYE) with reference to DDB1. (B) Cartoon illustration of SALL4A/B-CRBN-DDB1-CUL4A-RbX1 protein interaction network. (C) Superimposed 3D structures of the SALL4A ZnF3-CRBN-DDB1(green) complex (PDB ID: 6UML) and DDB1(yellow)-CUL4A-Rbx1-ZFC 2-4 of SALL4A complex (generated by protein-protein docking) with reference to DDB1 domain.

**Figure EV5.**
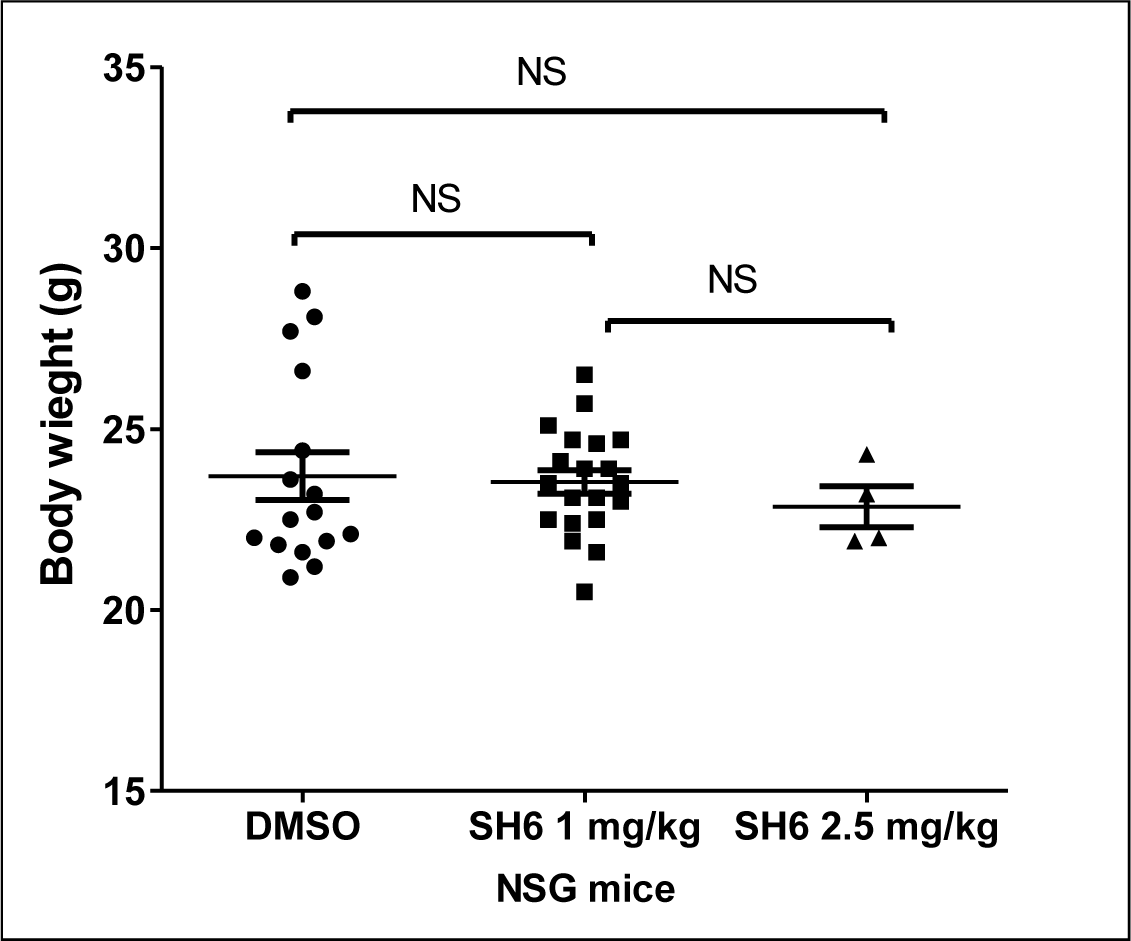
Body weight of mice at end point from three treatment groups (DMSO, SH6 1mg/kg, SH6 2.5mg/kg) demonstrates no significant difference between the groups.

**Table EV1.**
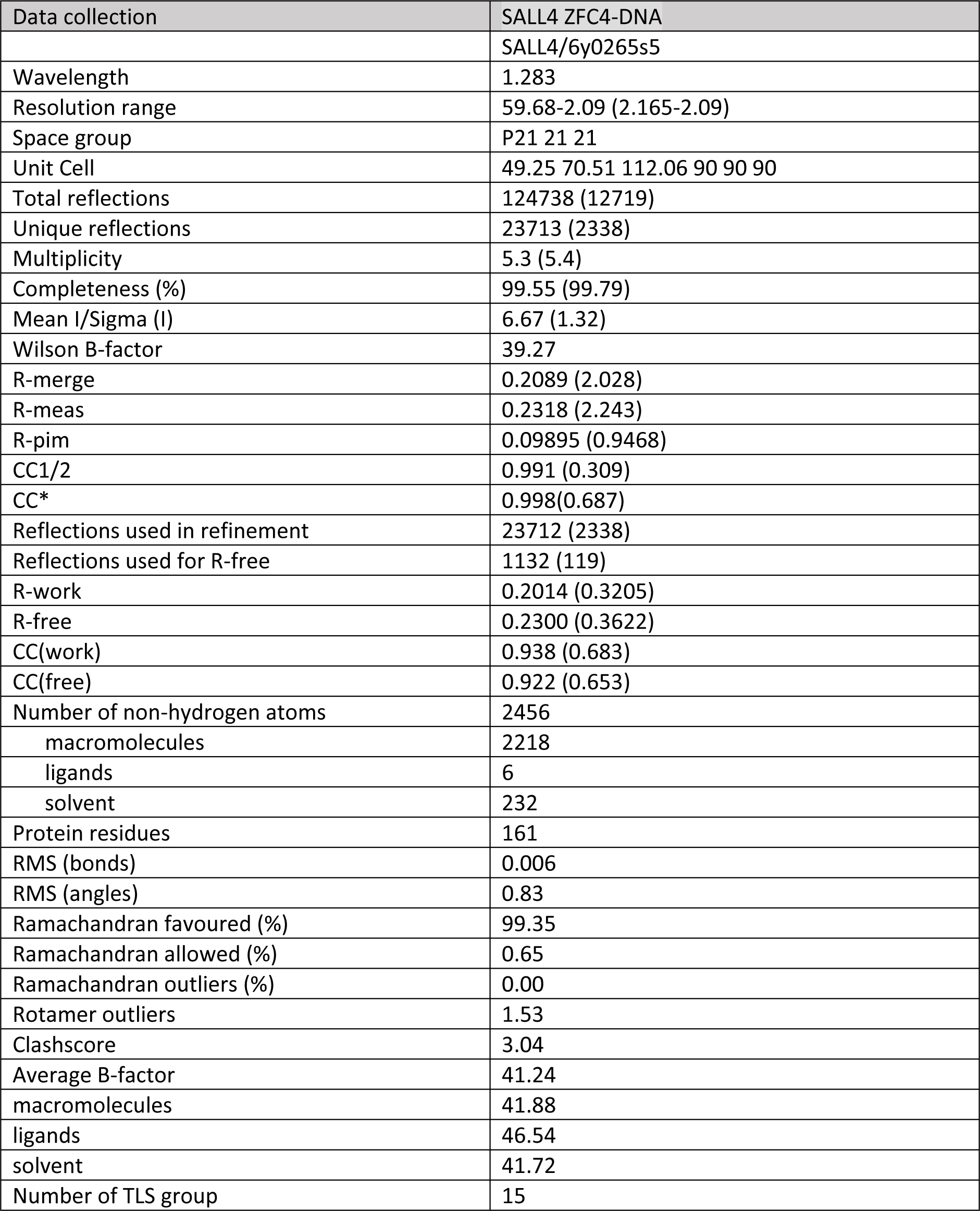
Crystallographic and refinement statistics of the ZFC4-DNA complex.

**Table EV2.**
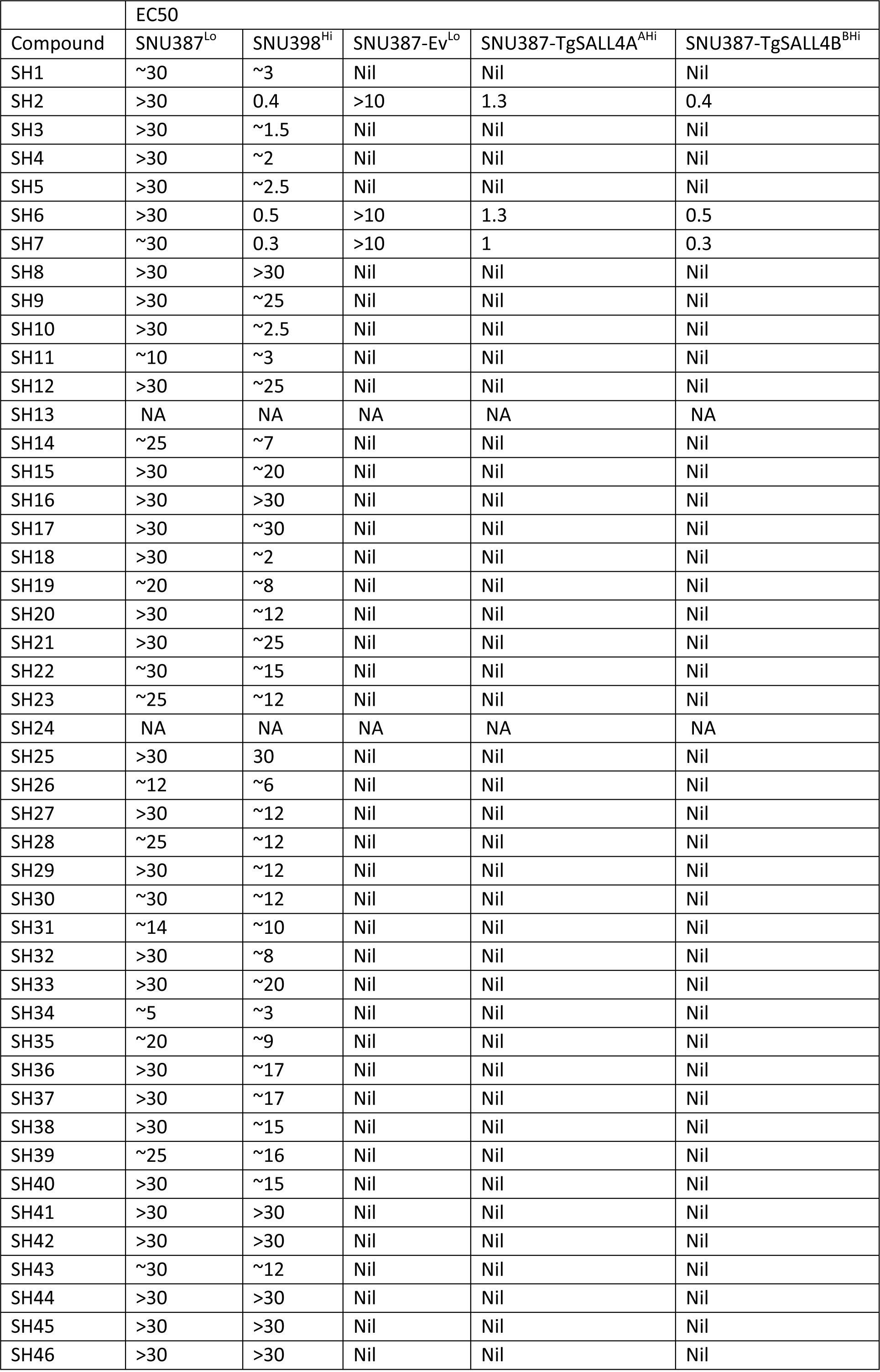
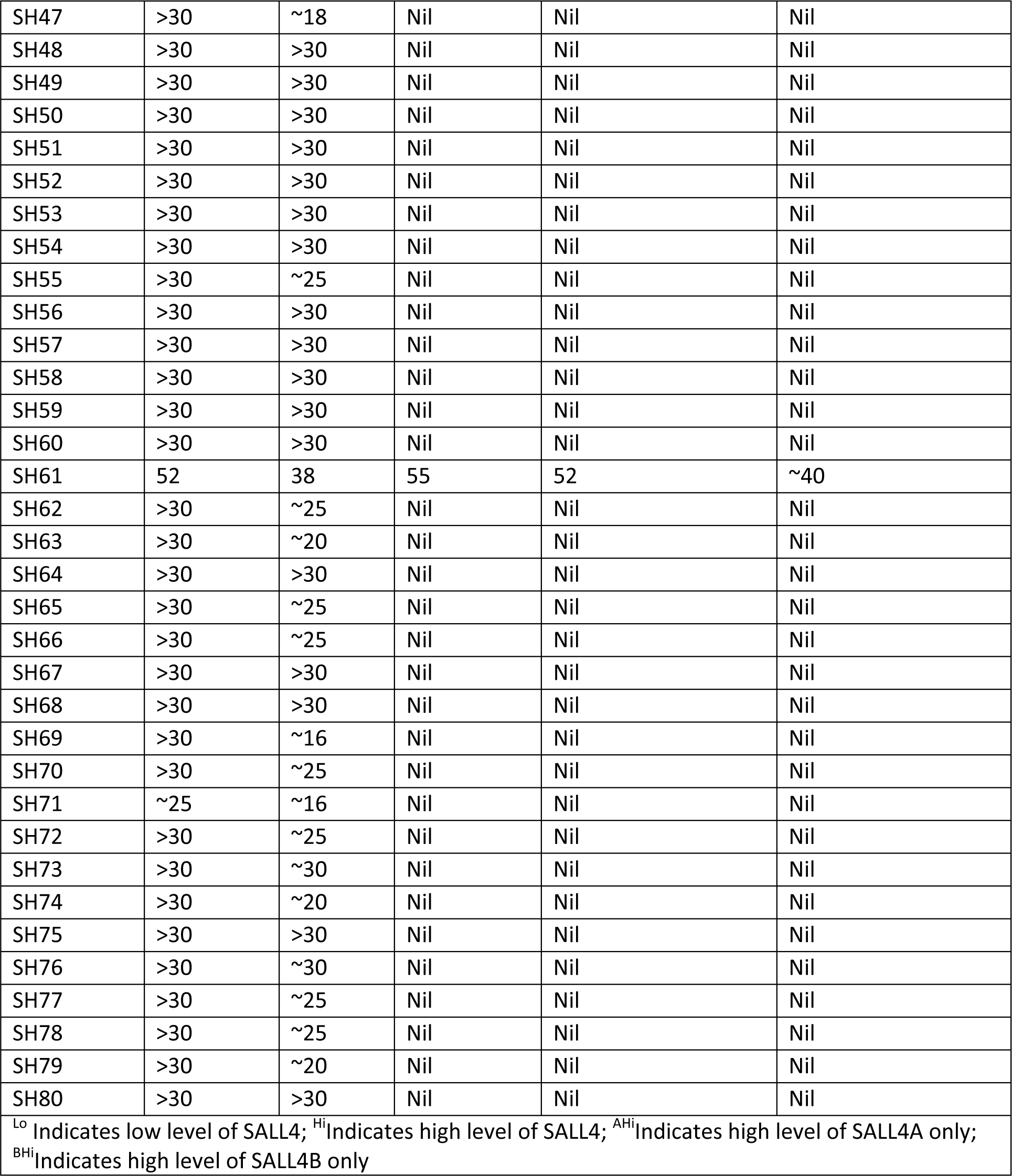
EC50 of cell lines treated with varying concentration of SH compounds.

**Table EV3.** Docking scores of SH6, SH61, and 3 thalidomide analogues against the predicted binding site in the SALL4A-CUL4A and the SALL4B-CUL4A complexes.

|  |  |  |
| --- | --- | --- |
| Compound | 50 | Docking scores |

|  | SALL4A-CUL4A | SALL4B-CUL4A |
| --- | --- | --- |
| SH6 | -10.12 | -10.21 |
| SH61 | -5.25 | -6.01 |
| Lenalidomide | -5.98 | -6.12 |
| Thalidomide | -7.01 | -7.45 |
| Pomalidomide | -6.26 | -6.39 |
| ZFC2-4 of SALL4A was docked to CUL4A and formed SALL4A-CUL4A complex. ZFC4 of SALL4B was docked to CUL4A and formed SALL4B-CUL4A complex. The same binding site was identified in both complexes and targeted in molecular docking of the five compounds. |  |  |

